# Identification and Functional Annotation of Long Intergenic Non-coding RNAs in the Brassicaceae

**DOI:** 10.1101/2021.09.17.460835

**Authors:** Kyle Palos, Anna C. Nelson Dittrich, Li’ang Yu, Jordan R. Brock, Larry Wu, Ewelina Sokolowska, Aleksandra Skirycz, Polly Hsu, Eric Lyons, Mark A. Beilstein, Andrew D. L. Nelson

## Abstract

Long intergenic noncoding RNAs (lincRNAs) are a large yet enigmatic class of eukaryotic transcripts with critical biological functions. Despite the wealth of RNA-seq data available, lincRNA identification lags in the plant lineage. In addition, there is a need for a harmonized identification and annotation effort to enable cross-species functional and genomic comparisons. In this study we processed >24 Tbp of RNA-seq data from >16,000 experiments to identify ~130,000 lincRNAs in four Brassicaceae: *Arabidopsis thaliana*, *Camelina sativa, Brassica rapa*, and *Eutrema salsugineum*. We used Nanopore RNA-seq, transcriptome-wide structural information, peptide data, and epigenomic data to characterize these lincRNAs and identify functional motifs. We then used comparative genomic and transcriptomic approaches to highlight lincRNAs in our dataset with sequence or transcriptional evolutionary conservation, including lincRNAs transcribed adjacent to orthologous genes that display little sequence similarity and likely function as transcriptional regulators. Finally, we used guilt-by-association techniques to further classify these lincRNAs according to putative function. LincRNAs with Brassicaceae-conserved putative miRNA binding motifs, short ORFs, and whose expression is modulated by abiotic stress are a few of the annotations that will prioritize and guide future functional analyses.

## Introduction

As genomic and transcriptomic analyses have become more prevalent, it has become clear that genomes are not solely composed of protein-coding genes, housekeeping RNAs, and transposable elements. One particularly important set of findings came from the Human ENCODE (ENCODE Project Consortium 2012) project where it was discovered that over 60% of the human genome is transcribed at some point in development into long non-coding RNAs (lncRNAs). The term “lncRNA” is a catchall for a class of transcripts united by two key features: a length > 200 nt and poor protein coding potential. The term lncRNA itself can be subdivided into natural antisense transcripts (NAT-lncRNAs), intergenic (lincRNAs), sense overlapping transcripts (SOT-lncRNAs), and intronic (int-lncRNAs). Each of these classes of lncRNAs appeared in analyses of RNA-seq data because they look like mRNAs (e.g., they are capped, polyadenylated, and often multi-exonic) (Guttman et al., 2009). However, most lncRNAs were missed or ignored in earlier EST based screens because of their low or tissue-specific expression and lack of open reading frames. RNA sequencing data from over 37,000 experiments reflecting 60 different tissues under different experimental and developmental conditions led to the identification of > 100,000 high confidence lncRNAs in humans (Volders et al., 2013; Zhao et al., 2019; Zhao et al., 2020).

In contrast to proteins, which were the focus of study long before the genomes from which they are encoded were sequenced, an appreciation for the abundance and varied roles of lncRNAs has primarily emerged along with the accumulation of sequencing data. As a result, the catalog of *functionally characterized* lncRNAs is limited, both in number and in diversity of organisms where they have been annotated (Statello et al., 2020; Seifuddin et al., 2020; Chekanova JA, 2021). Moreover, the extent to which functionally characterized lncRNAs are archetypal across plants, animals, and fungi is unknown. Thus, lncRNA identification and functional characterization lags far behind similar efforts in proteins, representing a fundamental gap in our understanding of how genomes operate.

Findings from across eukaryotes serve to illustrate the importance of lncRNAs to genome stability and regulation. Prominent mammalian examples include the telomerase RNA component (TERC), a scaffolding RNA that is crucial for chromosome maintenance (Feng et al. 1995); XIST, a guide RNA responsible for X chromosome inactivation (Brown et al. 1992); and HOTAIR, a developmentally-linked signaling RNA (31). In Arabidopsis, TERC has been characterized, with sequence and structural homologs present across the plant lineage, highlighting the potential for lncRNA conservation over long evolutionary timescales (Dew-Budd et al. 2020); (Fajkus et al. 2019); (Song et al. 2019). Most other lncRNAs functionally characterized in plants, such as COOLAIR, ELENA1, SVALKA, MAS, APOLO, and HID1, change expression or function in response to environmental cues, and can thus be classified as environmental sensors (Csorba et al. 2014); (Seo et al. 2017); (Kindgren et al. 2019); (Zhao et al. 2018; Ariel et al. 2020; Y. Wang et al. 2014). These examples reflect the myriad of different mechanisms by which lncRNAs play important biological roles in plants, and also likely represent the tip of the iceberg of what remains to be discovered.

One critical factor behind the dearth of functionally described lncRNAs in plants relative to mammalian systems is the lack of annotated lncRNAs, and, where lncRNAs have been annotated, the disparity in criteria and transcriptional data used for identification. In Arabidopsis, the bulk of annotated lincRNAs are derived from two studies (Amor et al. 2009); (Liu et al. 2012), although other genome-wide examinations have been performed (Moghe et al. 2013); (Y. Wang et al. 2014). The former examined full length cDNA libraries for lack of coding potential, whereas the latter utilized TILING arrays to infer gene structure and transcriptional status. In both cases the maximum allowable ORF was 100 AA or less. Other lincRNA identification efforts outside of Arabidopsis (e.g., GREENC), used official genome annotations generated by MAKER (Cantarel et al. 2008) without direct transcriptional evidence and maximum allowable ORFs of 120 AA. Yet in other systems, lincRNA identification efforts are limited to a few tissues or developmental stages (Qi et al. 2013; Moghe et al. 2013; L. Li et al. 2014; Shuai et al. 2014). While functional lncRNAs have been identified from many of these efforts, the disparity in identification schemes, and discordant developmental stages and environmental conditions makes it difficult to make sequence or structural based comparisons within and across species as is typically done for protein-coding genes.

Here we present a comprehensive and unified annotation of lincRNAs, using criteria established in mammalian systems, across four model or agriculturally significant Brassicaceae: *Arabidopsis thaliana*, *Camelina sativa*, *Brassica rapa*, and *Eutrema salsugineum*. We reprocessed over 16,000 different publicly available RNA-seq experiments (> 24 Tbp of raw data), and created our own Oxford Nanopore (ONT) and Illumina RNA-seq data, to identify lncRNAs in each of these species. We focus primarily on the intergenic class of lncRNAs for evolutionary and technical reasons: the evolution of NAT and SOT-lncRNAs is obscured by the overlapping protein-coding gene, and the unstranded nature of much of the publicly available RNA-seq data makes confident strand assignation of single exon transcripts difficult. Using transcriptomic, proteomic, epigenetic, and genome-wide RNA-protein interaction data, we examined our lincRNA catalog for features that separate and define lincRNAs from other transcriptional units. We used evolutionary and comparative genomic approaches, leveraging the unique strength of plant polyploidy, to identify conserved lincRNAs among the four species and the rest of the Brassicaceae as well as identify conserved motifs for functional testing. Finally, we used all of these contextual clues, as well as guilt-by-association techniques, to assign putative function to lincRNAs within our catalog.

## Results

### Identification of lincRNAs within the Brassicaceae

To overcome difficulties in comprehensive lincRNA identification (low-expression, tissue/environmental specificity) in our focal species (*Arabidopsis thaliana*, *Brassica rapa*, *Camelina sativa*, and *Eutrema salsugineum*), we processed all RNA-seq data deposited to the Sequence Read Archive (SRA) at the NCBI [accessed December, 2018] for these species. We excluded SRAs with epigenetic mutants, degradome experiments (GMUCT and PARE), small RNA-sequencing, and experiments with low sequencing depth (fewer than 1 million quantified/mapped reads; **Figure 1**). In addition to publicly available short read RNA-seq, we also performed Oxford Nanopore Technology (ONT) PCR-free cDNA sequencing on three tissues (10-day seedlings, 4-week mature rosettes, and open flowers) for the four focal Brassicaceae. We used previously developed workflows (Peri et al. 2019) utilizing the CyVerse computational infrastructure (Merchant et al. 2016) to map, in high throughput, ~24 terabases of RNA-sequencing data associated with 16,076 experiments (listed in **Supplemental File 1**) to their respective genomes. We then identified putative lncRNAs using the Evolinc computational pipeline (Nelson et al. 2017). After identification, we proceeded to filter the initial candidate lincRNAs based on a set of hierarchical filters similar to those used to identify the “gold standard” set of human lincRNAs by Cabili et al, 2011.

**Figure 1:**
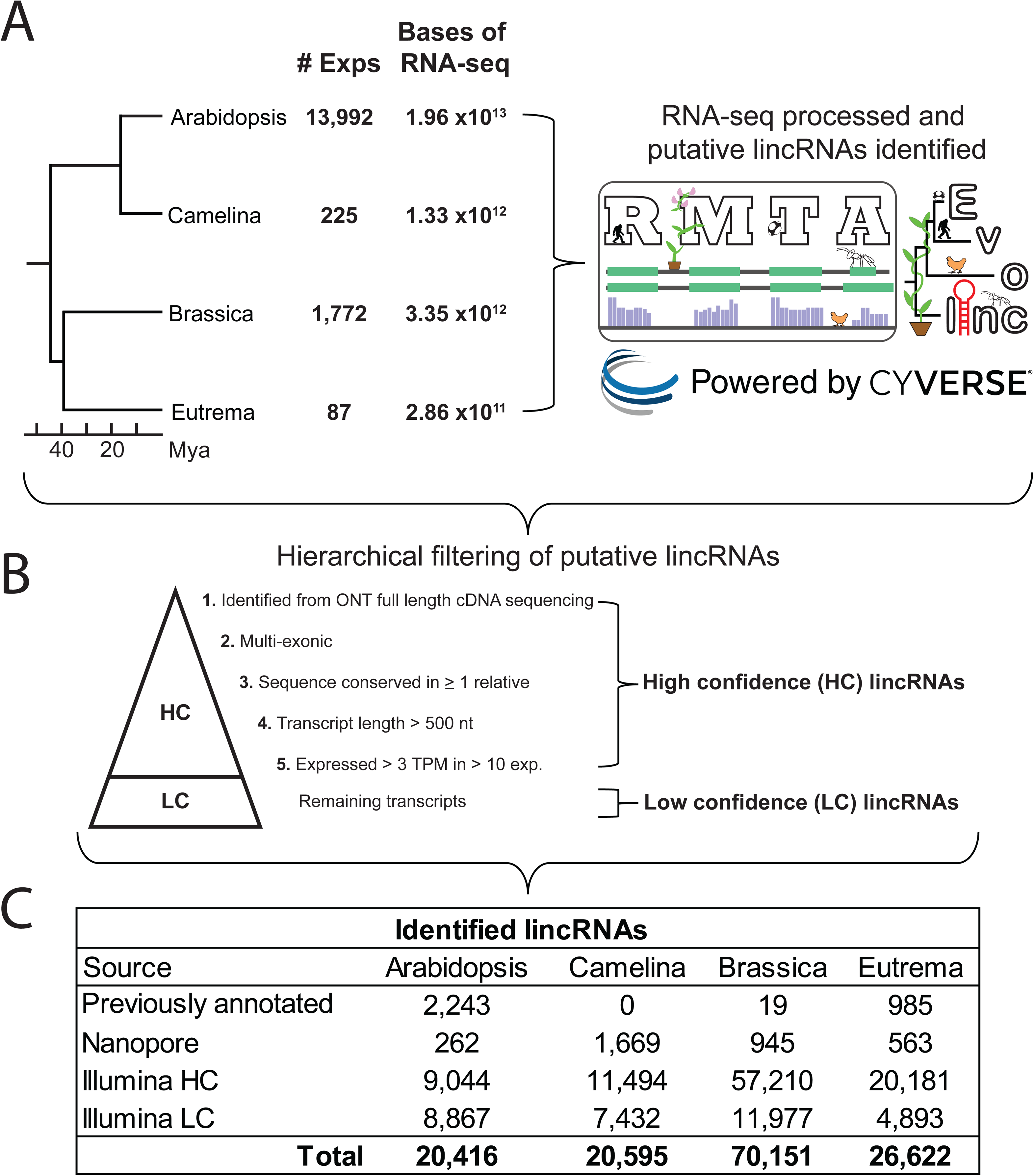
Basic identification and characterization of lincRNAs in each of the four target Brassicaceae. **A**) Number of experiments processed for each experiment using the RMTA and Evolinc pipelines in CyVerse’s cloud computing infrastructure. **B**) The metrics used for lincRNA additional hierarchical filtering. Note, lincRNAs only had to pass one additional filter to be considered a high confidence lincRNA. **C**) The number of identified lincRNAs.

The first of these filters pertained to whether we observed these lincRNAs in our ONT sequencing data. Due to the potential for full-length coverage of the ONT cDNA sequencing (Seki et al. 2019), any lincRNAs identified by ONT cDNA sequencing were annotated as high confidence (HC) without further filtering. Using ONT cDNA sequencing we identified 200 unannotated (i.e., not present in the Araport 11 annotation) lincRNAs in *Arabidopsis thaliana*, 945 in *Brassica rapa*, 1,669 in *Camelina sativa*, and 563 in *Eutrema salsugineum*. Due to concerns with transcriptional artifacts common in short read RNA-seq at lowly expressed loci, we next retained lincRNAs as HC if they were multi-exonic. This filter selects for transcripts that are less likely to be artifacts of transcript assembly algorithms (Cabili et al. 2011). By this criterion, 678, 12,422, 6,200, and 1,812 multi-exonic lincRNAs were identified and annotated as HC in Arabidopsis, Brassica, Camelina, and Eutrema, respectively (**Figure 1C**). Cognizant of previously identified and characterized mono-exonic lincRNAs (Lorenzi et al. 2021), (Sun and Ma 2019), (West et al. 2014)), we next filtered the remaining lincRNAs (mono-exonic) and labeled them as HC if 1) the transcript was conserved at the sequence and synteny level in the genome of at least one other Brassicaceae (See Materials and Methods), 2) the transcript length was > 500 nts, or 3) the transcript was expressed above 3 transcripts per million (TPM) in at least 10 RNA-seq experiments. All putative Evolinc lincRNAs that did not pass these filters were retained within our dataset as low confidence (LC) lincRNAs, as there is the risk that these transcripts represent false positives in our dataset. See **Supplemental File 2** for a complete list of HC and LC lincRNAs for each species. Thus, we identified a total of 9,244, 58,155, 13,163, 20,744 high confidence lincRNAs (HClincRNAs) in Arabidopsis, Brassica, Camelina, and Eutrema, respectively (**Figure 1B**). In contrast, 8,867, 11,977, 7,432, 4,893 low confidence lincRNAs were identified in the respective species.

Another source of false positives that we hoped to address comes from mis-annotating a transcript as a lncRNA when it in fact represents a misassembled or fragmented mRNA or is instead an extension of an annotated gene (e.g., a 5’UTR extension). To determine the frequency at which we were recovering false positive lincRNA assignments, we compared independently assembled transcriptomes from Illumina short read and ONT long read derived lncRNA datasets, searching for short read derived “lncRNAs” that were instead 3’ or 5 extensions of an adjacent gene within our ONT sequencing data. Using this approach we identified 39 lincRNAs in Arabidopsis that shared at least 1 ONT sequencing read with a neighboring mRNA (must be on the same strand) out of 2,370 lincRNAs for which we obtained ONT coverage ≥ 1. Of the 39 lincRNAs with overlapping sequencing reads, only 2 appeared to be bonafide mRNA extensions (**Supplemental Figure 1A**). The 37 other lincRNAs appear to share sequencing reads due to mis-assemblies or genomic DNA contamination in the sequencing (**Supplemental Figures 1B and 1C**, asterisks), or are larger variants of Araport lncRNAs. In general we identified strong agreement between ONT and Illumina derived lincRNA transcript models (**Supplemental Figure 1D**), suggesting the depth of Illumina sequencing used here was more than sufficient to overcome misassembly common for lowly expressed transcripts. Given the low rate (1.64%) of false positives, we remain confident that the transcripts we have identified are indeed independently transcribed elements within the Arabidopsis genome.

We next assessed how many of the previously identified Arabidopsis lncRNAs were expressed in our assembled RNA-seq data. Given the comprehensive nature of our dataset, we presumed that a prior annotated lncRNA was misannotated if we did not observe expression above 1 TPM in at least 10 Arabidopsis RNA-seq datasets. There are ~ 3,000 annotated lncRNAs within the Araport11 genome annotation, a group that includes 2455 “lnc_RNAs”, 286 “ncRNAs”, and 726 “novel transcribed regions”. To create a uniform dataset of lincRNAs, we filtered out transcripts that did not fit the most basic definitions of a lincRNA (over 200 nt and not overlapping a protein coding gene), and for which we did not observe expression. Of the 2455 “lnc_RNAs”, 401 were removed for protein-coding gene overlap. A further 157 were relabeled as poor support due to lack of sufficient expression levels based on our expression filtering mentioned above (> 1 TPM in 10 experiments). However, we did observe low levels of expression (> 0.1 TPM) for some of these “poorly supported” genes in various tissue expression atlases, stress datasets, or our Nanopore sequencing (**Table 1**). In total, we confirmed 1,897 Araport lnc_RNAs to be HC-lincRNAs. For the 286 annotated “ncRNAs”, 189 (66%) passed the length, intergenic, and expression criteria. Finally, we analyzed the novel transcribed regions and first assessed coding capacity as these features were not originally annotated as noncoding. We treated these transcripts to the same set of filters as our lincRNA dataset (ORF < 100 AA, longer than 200 nts, poor coding potential), which resulted in 571 NTRs annotated as lincRNAs and included in further analyses. In total, we reclassified 2,566 Araport genes as lincRNAs (**Supplemental File 2**). Details on the lnc_RNAs, ncRNAs, and NTRs that were removed from the final dataset can be found in **Supplemental File 7**.

**Table 1:** Broad categories of experiment/tissue in which highly context-specific lincRNAs were found to be expressed in Arabidopsis and Brassica.

| Broad Category | Arabidopsis | Brassica |
| --- | --- | --- |
| Embryo-associated | 468 | 143 |
| Dissected flower tissue | 648 | 77 |
| Biotic infection | 215 | NA |
| Epigenetic mutants | 47 | 15 |
| Root tip or meristem | 3,448 | NA |
| Shoot meristem | 1,788 | NA |
| Mixed accessions/ssp. | 363 | 21,840 |
| RILs | NA | 19,097 |
| Genetic mutants | 2,089 | NA |
| Circseq | 452 | NA |
| Other | 2,773 | 2,202 |
| **Total** | **12,291** | **43,374** |

The definition used by the community to identify lncRNAs is arbitrarily set and may include transcripts that encode for small proteins. Prior results have demonstrated previously annotated lncRNAs as bound to ribosomes, and in some cases have identified protein products for certain “lncRNAs” (Ji et al. 2015; Hsu et al. 2016; Wu et al. 2019). We used Ribo-seq (Ingolia et al. 2009; Wu and Hsu 2021) and protein mass spectrometry (MS; (Domon and Aebersold 2006)) data from Arabidopsis seedlings (PRIDE: PXD026713) to identify translated short ORFs (sORFs) and protein products within our “lincRNAs’’. Out of the 1,172 lncRNAs expressed > 0.1 TPM in these experiments, we uncovered evidence of translation for 158 lncRNAs (120 Ribo-seq/38 MS) ranging in size from 3-136 amino acids (**Figure 2A**). There is no correlation between lincRNA and sORF length (**Supplemental Figure 2**), but we did observe a tendency for previously identified lincRNAs (i.e., Araport-derived, n = 81) to contain longer sORFs than Evolinc-derived lincRNAs (n = 77; p-value 0.046; **Figure 2A**), likely due to the more restrictive criteria used to annotate the Evolinc lincRNAs. LincRNAs containing sORFs have been denoted as such in **Supplemental File 2**, but as they reflect previously unidentified genes that would otherwise have been called lincRNAs, were retained as a separate set of transcripts for downstream analyses.

**Figure 2:**
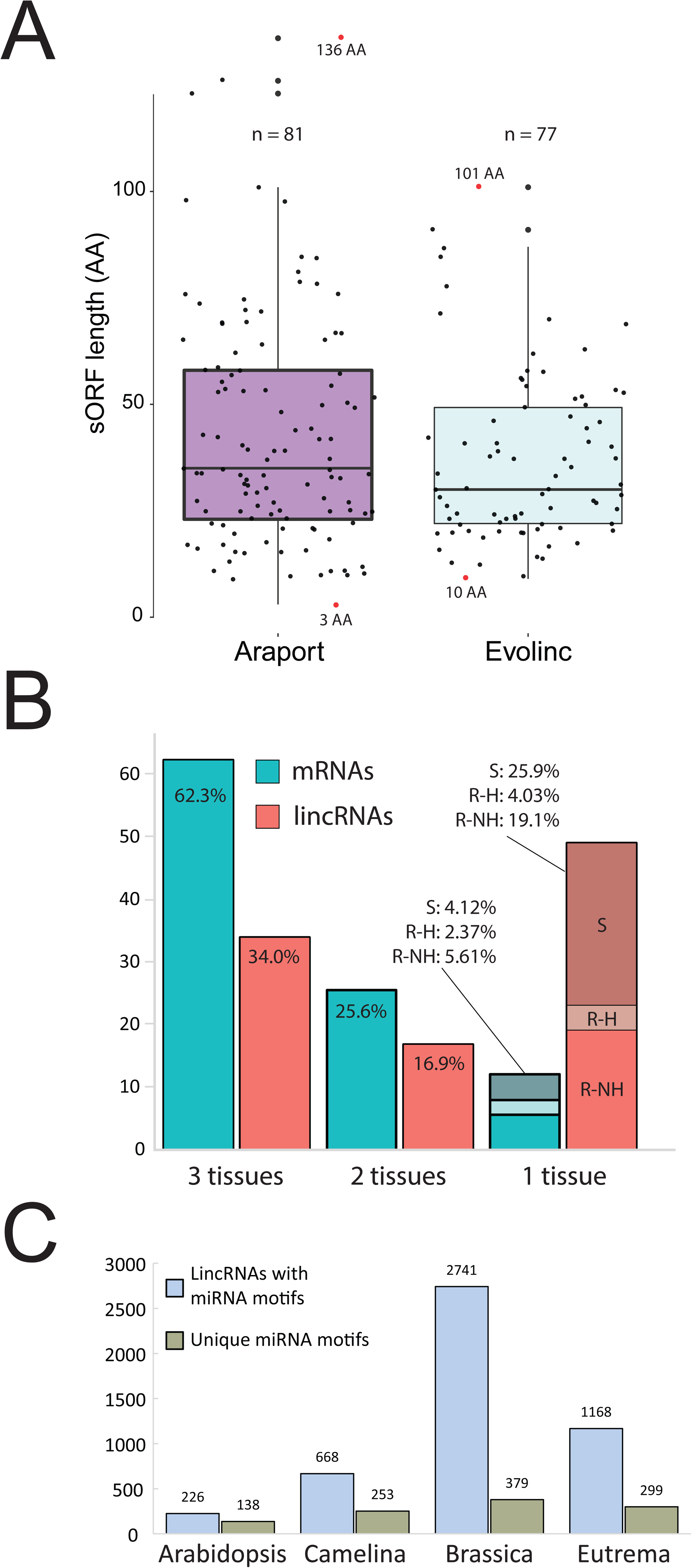
Identification of functional motifs within Arabidopsis lincRNAs. **A**) Distribution of the length of the newly identified sORFs within the Araport (previously identified) and Evolinc (this study) lincRNA populations. The largest and smallest sORFs are labeled (red dot) with the length denoted (in amino acids, AA). **B**) Distribution of identified structured and protein-bound elements within the total Arabidopsis lincRNA dataset based on PIP-seq data from three different tissues (S = Seedling, R-H = Root with hairs, R-NH = Root with no hairs; see **CITE** for more details). The percent of unique mRNAs (blue scale) or lincRNAs (red scale) that overlapped with at least one PIP-seq read are shown. **C**) Frequency of miRNA binding sites within lincRNA populations from each of the four focal species. Total number of lincRNAs in each of the four species containing a putative miRNA binding site is shown in blue, with unique miRNA motifs shown in grey/green.

We next used publicly available transcriptome-wide protein-interaction profile sequencing (PIP-seq, (Foley et al. 2017; Gosai et al. 2015)) data from roots (hair and nonhair) and seedlings (GEOs GSE58974 and GSE86459) to identify lincRNAs in our dataset that were protein bound and for which we have some measure of structure. Within the three tissues, we identified 397 structured and protein-bound lincRNAs. 135 (34%) lincRNAs were identified in all three datasets, whereas 195 were restricted to a single tissue (**Figure 2B**). Of these tissue-restricted lincRNAs, 119 were found to be structured in root cells, with the vast majority (103; 26% of structured lincRNAs) only present in non-hair root cells (R-NH; **Figure 2B**). In contrast, most mRNAs (62%) were found to be structured in all three tissues, whereas only 6% were restricted to non-hair root cells (**Figure 2B**). Thus, we have evidence for a subset of the Arabidopsis lincRNAs that are protein-bound and structured, with some evidence of tissue-specificity. These lincRNAs have been annotated in **Supplemental File 2** and the MSAs are available in the CyVerse Data Store (DOI).

Some lincRNAs are known to interact with miRNAs, either in a competitive inhibitory fashion (i.e., miRNA sponge; (Zhang et al. 2019), or to directly regulate the lincRNA itself (e.g., TAS1A; (Howell et al. 2007; Chen, Li, and Wu 2007)). Using the miRNA binding site prediction tool psRNATarget (Dai, Zhuang, and Zhao 2018) we identified 226 Arabidopsis lincRNAs with at least one putative miRNA recognition site (**Figure 2C**). Importantly, within this set of lincRNAs we identified previously identified miRNA-regulated lincRNAs such as TAS1A and TAS1B. We identified a further 668 miRNA-interacting lincRNAs in Camelina, 2,741 in Brassica, and 1,168 in Eutrema (**Figure 2C**). These putative miRNA-interacting lincRNAs, with their interaction sites, are annotated in **Supplemental File 5**. In sum, we used a wealth of public information to improve the genome annotations of four agricultural or model Brassicaceae.

### Fundamental features of Brassicaceae lincRNAs

We next examined basic characteristics of our lincRNA datasets with the goal of identifying features that might improve future lncRNA identification efforts. LincRNAs in all four species have significantly lower GC content relative to protein-coding genes (*P* value for all species’ lincRNA-mRNA comparison < 2.2e-16, Wilcoxon Signed Rank test; **Figure 3A**). Additionally, transcript length of lincRNAs are significantly shorter than mRNAs (*P* value for all species’ lincRNA-mRNA comparison < 2.2e-16, Wilcoxon Signed Rank test; **Figure 3B**). Interestingly, when we looked at multi-exonic lincRNAs and mRNAs, we found that the average length of individual exons of lincRNAs are significantly longer than the average length of individual exons of mRNAs for all species except *Camelina sativa,* where lincRNA exons displayed a similar trend to the other species (*P* value for all species < 2.2e-16, Wilcoxon Signed Rank test; **Figure 3C**). Finally, we analyzed the distribution of exons in lincRNAs in all four species. LincRNAs in Arabidopsis are mostly mono-exonic (~ 91.1 %), while the lincRNAs identified in the other species have a much more balanced distribution of exon counts, but still have fewer exons than mRNAs on average (**Supplemental Figure 3A**).

**Figure 3:**
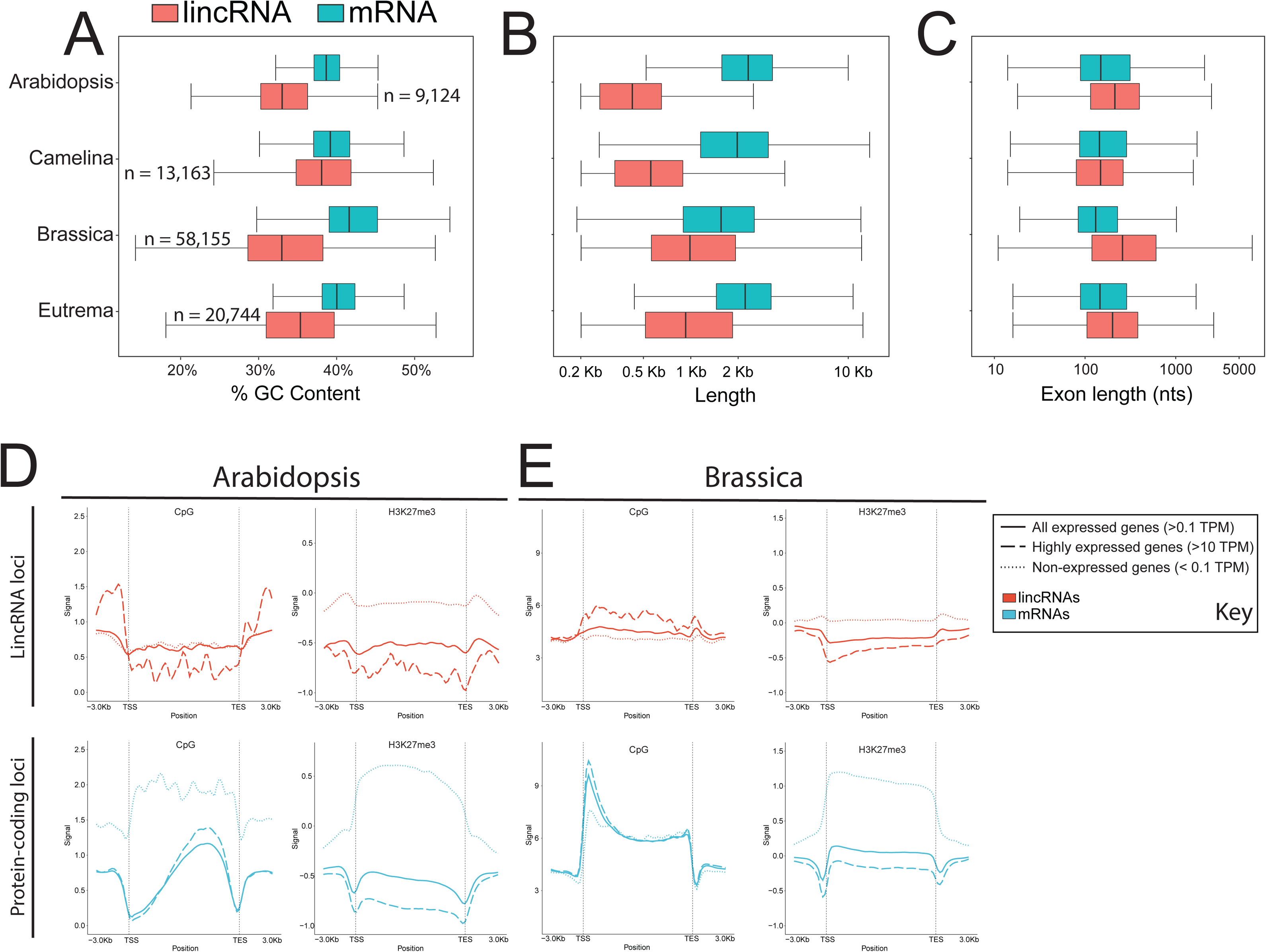
Basic sequence characteristics of Brassicaceae lincRNAs. **A**) % GC content and **B**) transcript length comparisons of mRNAs and lincRNAs in each of our 4 focal Brassicaceae. **C**) Exon length distribution for mRNAs and lincRNAs. All comparisons are significant (p < 2e-16) based on a Wilcox test with Bonferroni correction. CpG DNA methylation and H3K27me3 epigenetic profiles for lincRNAs and mRNAs in **D**) Arabidopsis and **E**) Brassica. LincRNAs and mRNAs were separated based on expression levels using paired RNA-sequencing data.

We next searched for differences in epigenetic regulation between lincRNAs and mRNAs. Owing to the wealth of genome-wide epigenetic data in Arabidopsis and Brassica, we identified experiments in both species for making direct comparisons, including CpG DNA methylation and H3K27 trimethylation (H3K27me3; see methods; **Supplemental File 4**). We further divided our gene sets based on expression to better understand the interplay between expression and epigenetic regulation. In Arabidopsis, lincRNA loci are distinct from both transposable elements and protein-coding loci in that they have a consistent decrease in CpG methylation across the gene body (**Figure 3D; Supplemental Figure 3B**). This decrease was largely consistent regardless of expression. In contrast, expressed protein-coding loci show the characteristic dip in CpG methylation at the transcription start site (TSS) and increase near the transcription end site (TES). TEs show elevated CpG methylation across the gene body relative to their surrounding genomic regions. This trend is reversed in Brassica, where protein-coding loci displayed a peak in CpG methylation at the TSS and lncRNAs showed elevated CpG methylation across the gene body (**Figure 3E**). Expressed protein-coding and lincRNA loci showed decreased levels of H3K27me3 both across the gene body and relative to non-expressed loci (**Figure 3D**), a pattern that was recapitulated in Brassica (**Figure 3E**). We also examined H3K9 acetylation, for which data were only available for Arabidopsis (**Supplemental Figure 3B**). Expressed Arabidopsis protein-coding loci displayed the characteristic peak in acetylation at the TSS and dip at the TES, whereas lincRNA loci displayed an increase in acetylation across the gene body that was positively associated with expression. In general, lincRNAs in Arabidopsis and Brassica are distinguished from protein-coding loci and TEs in that they display similar patterns across their gene body to the patterns associated with the TSS of protein-coding loci, a feature that becomes more pronounced with higher expression.

LncRNAs in mammalian systems are often tissue or cell-type specific, and often lowly expressed at the tissue level relative to mRNAs. This has also been observed to a certain extent in plant systems as well, albeit with far fewer tissue comparisons. Maximum lincRNA expression, in any tissue, was indeed ~10-fold lower compared to mRNAs in all species examined (**Figure 4A and Supplemental 4A**). Tissue specificity (TAU; (Yanai et al. 2005)) was determined based on expressions data from tissue atlases in Arabidopsis ((Klepikova et al. 2016) and Brassica (Tong et al. 2013; Bilichak et al. 2015), as well as from our ONT RNA-seq data. As expected, lincRNAs from all four species were, on average, significantly more tissue-specific than their respective mRNA cohorts (**Figure 4B and Supplemental Figure 4B**). We also observed a negative correlation between lincRNA tissue specificity and expression, a feature that was significantly more pronounced than for mRNAs (**Figures 4C and 4D**). This negative correlation was observed across multiple tissues (e.g., female reproductive, leaf, and male reproductive; **Supplemental Figure 4C**), although we did observe tissue-dependent differences, such as high expression associated with high specificity for both lincRNAs and mRNAs in pollen/anther RNA-seq data. The sORF containing lincRNAs displayed expression and tissue specificity values similar to mRNAs (**Figures 4A and 4B**), further supporting an mRNA assignment. Given this link between lower tissue specificity and coding potential, we more closely examined the Arabidopsis lincRNAs (n = 89) with TAU values lower than the median value for mRNAs (TAU < 0.502; **Figure 4B**, black box). Based on sequence similarity, these broadly expressed lincRNAs do not appear to be recently pseudogenized protein-coding genes, but, for a subset (n = 61), expression is significantly correlated with a neighboring gene less than 500 bp away (**Supplemental Figure 4D**). Thus, high tissue specificity and low expression can be considered a defining feature of Brassicaceae lincRNAs and can potentially help to distinguish unannotated sORF containing transcripts.

**Figure 4:**
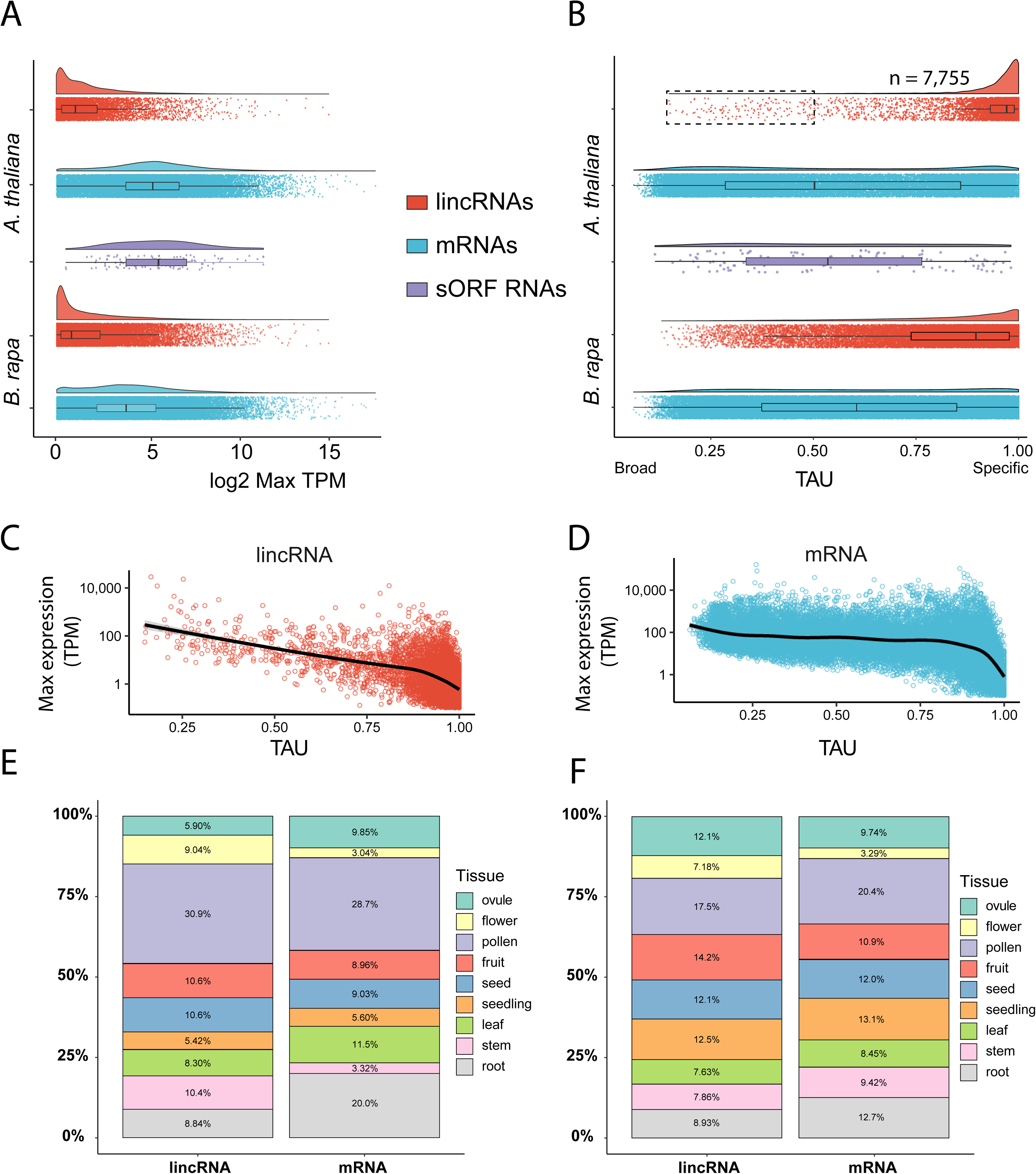
Expression dynamics of Arabidopsis and Brassica lincRNAs. **A**) Log2 max TPM for lincRNAs, mRNAs, and sORF containing lincRNAs (Arabidopsis only) using tissue atlas data for the two species. **B**) Tissue specificity (TAU) for Arabidopsis and Brassica transcripts. The dashed box denotes the 96 Arabidopsis lincRNAs that are below the median TAU value of mRNAs and were inspected further for similarity to protein-coding genes (see text for details). **C-D)** Correlation between tissue specificity and max expression for Arabidopsis lincRNAs (**C**) and mRNAs (**D**) within the Klepikova tissue atlas. **E-F**) Stacked bar plots describing where lincRNAs or mRNAs are most highly expressed in Arabidopsis (**E**) and Brassica (**F**).

In mammalian systems, a large number of lincRNAs are expressed, or show elevated expression, in male reproductive tissues ((Hong et al. 2018)). This phenomenon is attributed to relaxed epigenetic control within these tissues. We sought to determine if this was also a feature of plant lincRNAs by examining lincRNA expression within the Arabidopsis and Brassica tissues atlases. Approximately 45 and 35% of lincRNAs in Arabidopsis and Brassica, respectively, were most highly expressed in reproductive tissues, with pollen being the predominant source of maximum expression levels (**Figure 4E and 4F**). A similar percent of mRNAs showed peak expression in reproductive tissues in the two species, suggesting a general transcriptome-wide, instead of lincRNA-specific, phenomenon. However, lincRNAs restricted to pollen tissue were expressed significantly higher than lincRNAs restricted to other tissues (e.g., female reproductive versus leaf tissue; **Supplemental Figure 4C**, note scales). To aid in exploration of lincRNA and mRNA expression between tissues and experiments, these data have been uploaded to the appropriate BAR eFP Browser (Provart and Zhu 2003), and are explorable through an interactive Clustergrammer (Fernandez et al. 2017) Jupyter notebook binder found at https://github.com/Evolinc/Brassicaceae_lincRNAs (**Supplemental Figure 5**).

**Figure 5:**
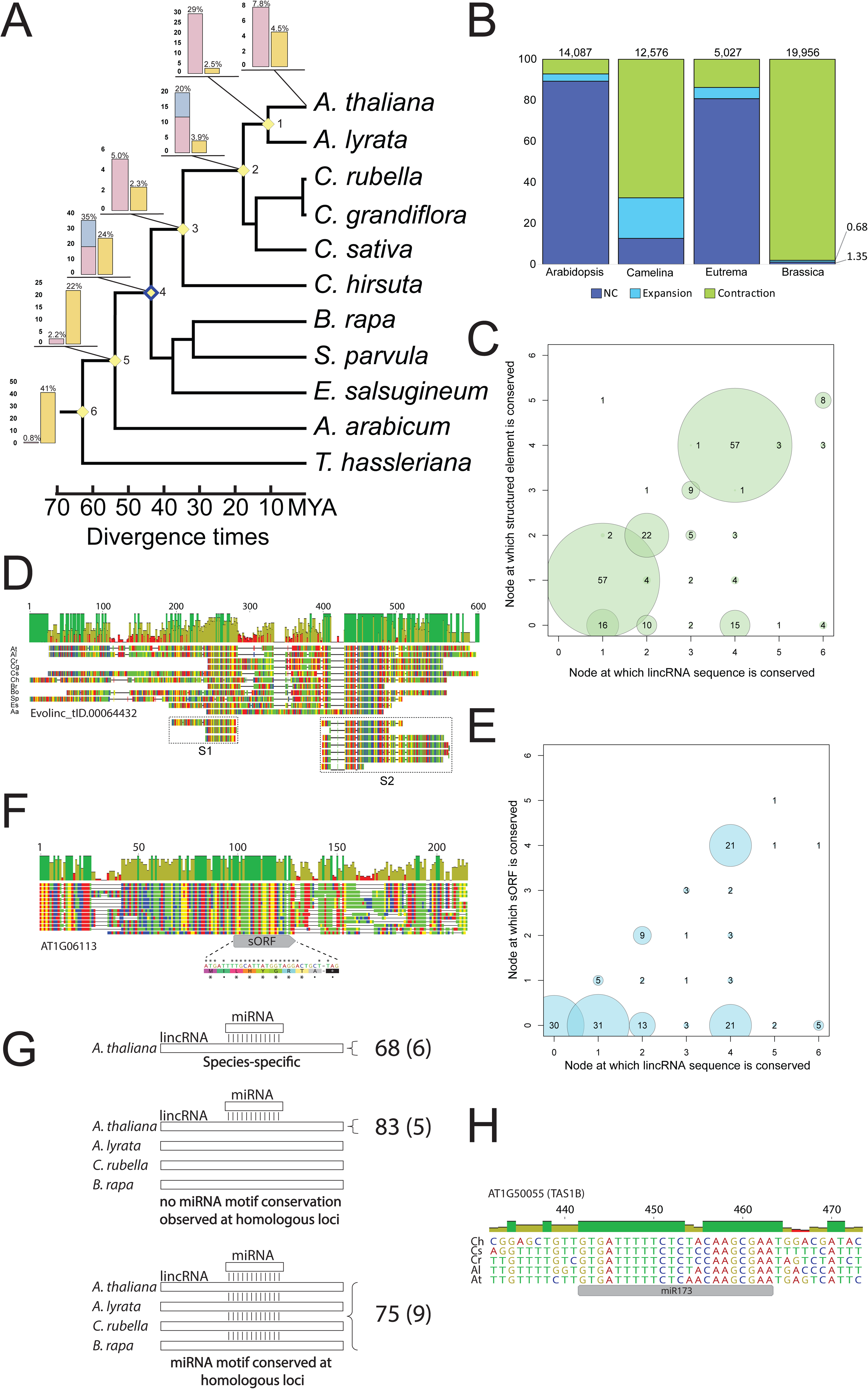
Sequence and transcriptomic conservation of Arabidopsis lincRNAs and their functional motifs across the Brassicales **A)** Arabidopsis lincRNA conservation across select Brassicale: *Arabidopsis thaliana*, *Arabidopsis lyrata*, *Capsella rubella, Capsella grandiflora, Camelina sativa*, *Cardamine hirsuta*, *Brassica rapa*, *Schrenkiella parvula*, *Eutrema salsugineum*, *Aethionema arabicum*, and *Tarenaya hassleriana* (representative of Cleomaceae). The inset bar graphs depict the percent of Arabidopsis lincRNAs and mRNAs (yellow bar) restricted to that node (out of 20,416 total lincRNAs and 27,173 mRNAs examined). For lincRNAs, pink bars represent lincRNA sequence homologs found at that node, whereas blue bars represent transcriptional syntelogs, and thus is dependent on lincRNAs having been identified in a species descending from that node. **B)** Percent of lincRNAs in each of the four focal species for which we could infer gene family expansion, contraction, or for which there was no change (NC) relative to at least 2 of the closest relatives for each species. See Methods for more information. **C**) Correlation between the nodes at which a lincRNA is conserved (from **A**) and at which the structured element (from Arabidopsis) is conserved. **D**) Multiple sequence alignment (MSA) of a structured and protein-bound Arabidopsis lincRNA (Evolinc_tID.00064432) where the functional motif and lincRNA are conserved to the same node. **E**) Correlation between lincRNA conservation and sORF conservation. **F**) MSA of a sORF containing Arabidopsis lincRNA (*AT1G06113*) where the lincRNA and sORF are conserved to the same node. **G**) Schematic demonstrating the number of putative miRNA binding motifs that were found to be either species-specific (lincRNA and miRNA motif are restricted to Arabidopsis), not conserved (lincRNA is conserved but conservation is not associated with miRNA motif), or conserved. The number in parentheses represents the number of lincRNAs with putative miRNA binding motifs that are also stress-responsive. **H**) Example MSA of a lincRNA (Arabidopsis *TAS1B*) with a conserved miRNA binding motif in Cardamine, Capsella, and Arabidopsis.

Interestingly, 48% and 60.8% of the complete (HC + LC) Arabidopsis and Brassica lincRNA datasets, respectively, were not expressed above 0.1 TPM in their respective tissue atlas suggesting these lincRNAs are not expressed under “normal” conditions during development. Considering that expression was a requirement for identification, we sought to determine where these “context-specific” lincRNAs (CS-lincRNAs) were expressed. We screened through all of the Arabidopsis and Brassica RNA-seq data looking for experiments of maximal expression. We extracted metadata from those experiments from the NCBI SRA and then grouped lincRNAs into similar categories based on expression (see **Materials and Methods**). In Arabidopsis, the majority of the CS-lincRNAs showed maximal expression in experiments that performed high-resolution sequencing of root or shoot meristems (n = 5236; Table 1), suggesting these lincRNAs are expressed in very limited cell types. 909 lincRNAs (~4.5% of dataset) were found to be expressed under stress (abiotic or biotic) conditions (**Table 1**). In Brassica, the vast majority of the CS-lincRNAs (n = 40,937; 57.5%) were maximally expressed in sequencing data sampling recombinant inbred lines (n = 19,097) or hybridization experiments with different Brassica accessions (n = 21,840; Cheng et al., 2016), indicating a high degree of transcriptional variation between genetic backgrounds. We also observed a subset of Arabidopsis lincRNAs (~350) that were only expressed in specific accessions or in crosses between accessions. Finally, 7,407 (10.4%) Brassica CS-lincRNAs were expressed under stress conditions. To allow researchers to sort lincRNAs based on their own priorities, expression metadata have been assigned to each CS-lincRNA in **Supplemental File 2**. These data highlight both the extreme tissue specificity possible for lincRNAs, as well as the potential for lincRNAs to be expressed during, and perhaps play a role in, growth and development of recent hybrids.

### Evolutionary features of Brassicaceae lincRNAs

As conservation is typically seen as a proxy for functional significance in protein-coding genes, we next sought to determine the degree to which lincRNAs from each of the four species were conserved across the Brassicaceae. Using each of the respective sets of lincRNAs as query, we first searched for sequence homologs within the genomes of nine Brassicaceae as well as *Tarenaya hassleriana*, a representative of the sister family Cleomaceae (Cheng et al. 2013). A comparative analysis of the complete Arabidopsis lincRNA dataset with Evolinc-II (Nelson et al. 2017) revealed that 32.9% (6,781) are species-specific (i.e., no sequence homologs are identified in any other species within the family; **Supplemental File 3**). Of the remaining ~14,000 lincRNAs, 6,045 (29% of total) are restricted to the genus Arabidopsis (node 1; **Figure 5A**). Much of these species or genera-specific conservation is driven by the low-confidence lincRNAs. The low-confidence lincRNAs were significantly more likely to be poorly conserved (nodes 0-1; **Supplemental Figure 6A**) than the high-confidence lincRNAs. Sequence homologs are present for ~35% (4,336) of the Arabidopsis lincRNAs at node 4, which represents the coalescence point between Brassicaceae lineages I and II (**Figure 5A**), suggesting these lincRNAs originated ≥ 43 MYA. The majority of these sequence homologs corresponded to either Evolinc-identified lincRNAs or unannotated intergenic sequence in each of the other species (**Supplemental Figure 6E**), suggesting that these lincRNAs have been evolving as lincRNAs, and not as pseudogenized loci, for the last 43 MY.

**Figure 6:**
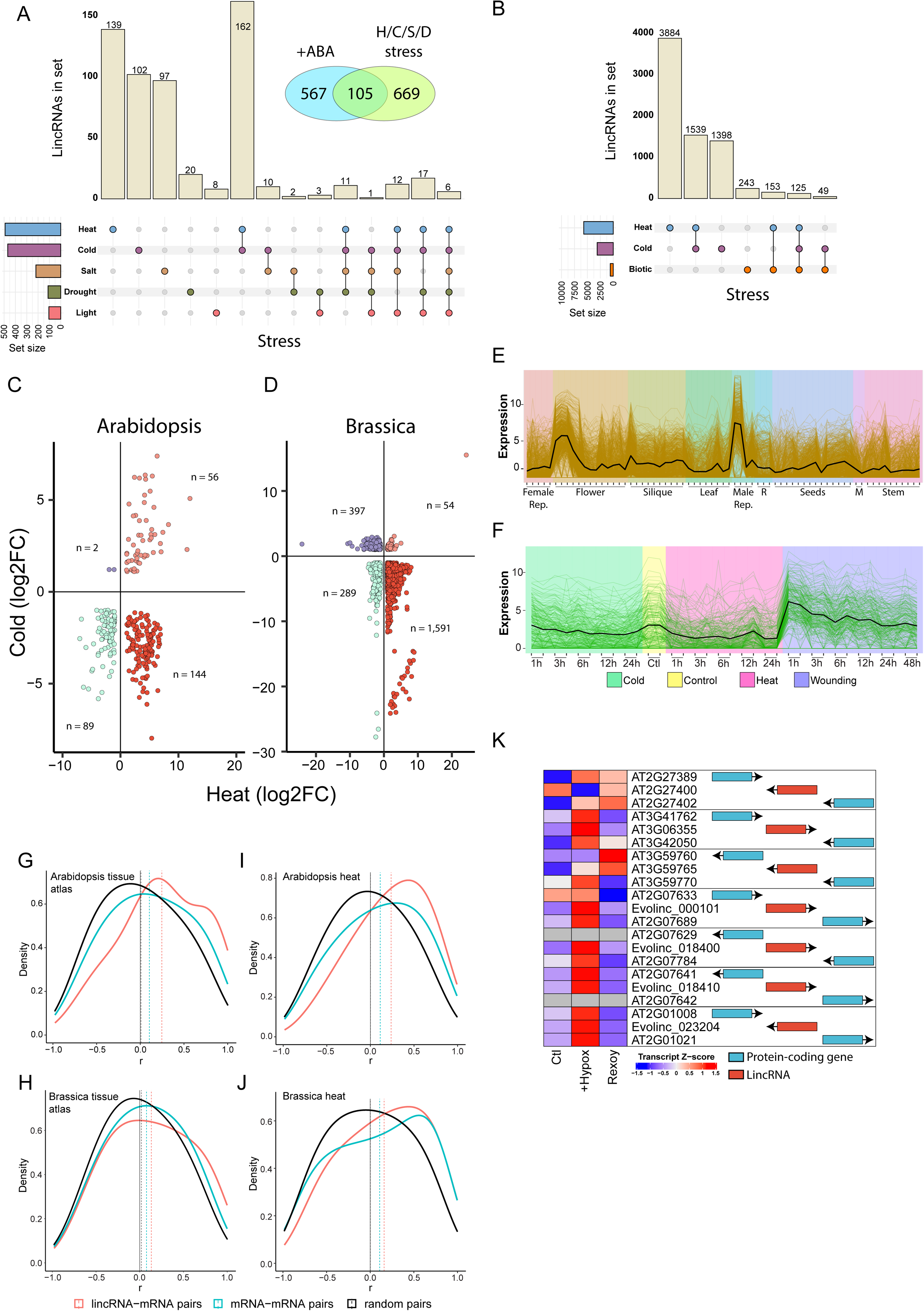
Inferring lincRNA function from transcriptomic data. **A-B**) Upset plots depicting the number of stress-responsive lincRNAs in Arabidopsis (**A**) and Brassica (**B**). The vertical tan bars depict the number of lincRNAs found in each stress, or combination of stresses, shown below. The horizontal colored bars depict the total number of lincRNAs associated with that stress across all combinations. For Arabidopsis, an inset Venn diagram depicts the number of lincRNAs found to be both stress (heat, cold, salt, or drought stresses) and ABA responsive. **C-D**) Scatterplots of temperature responsive lincRNAs in Arabidopsis (**C**) and Brassica (**D**). **E-F**) Modules of similarly expressed mRNAs and lincRNAs from the Arabidopsis Klepikova tissue atlas (**E**) or stress atlas (**F**). “M” = meristem tissue, “Male Rep.” = male reproductive tissues (stamens and anthers). **G-J**) Density plots showing the distribution of expression correlation between different gene pairs in Arabidopsis and Brassica tissue atlases (**G**-**H**) as well as Arabidopsis and Brassica heat stress experiments (**I-J**). **K**) Summary diagram of lincRNAs that are both hypoxia-stress responsive and bound by the HRE2 transcription factor. On the left are Z-transformed expression data for each lincRNA and their closest protein-coding gene. On the right is a depiction of the arrangement and orientation of each lincRNA and protein-coding gene set. **L**) Bar plot depicting the number of lincRNAs for which we have varying degrees of functional evidence.

The percent of species-specific lincRNAs for the other three species ranges from 49% (C. sativa) to 75% (E. salsugineum; **Supplemental Figures 6B-D**). Sequence homologs for ~0.8% (162) of the Arabidopsis lincRNAs were recovered in *T. hassleriana*, with similarly low percentages (3, 1.5, and 1) of sequence homologs identified for Camelina, Brassica, and Eutrema lincRNAs, respectively (**Supplemental Figures 6B-D**). In sharp contrast to lincRNAs, sequence homologs were recovered for > 43% of Arabidopsis protein-coding genes in *T. hassleriana* (**Figure 5A**). Thus, while a majority of Brassicaceae lincRNAs are species-specific, for each of the four focal species, a subset of lincRNAs display higher rates of sequence conservation across the family and likely depend on sequence for function. From these data we highlight several functionally characterized Arabidopsis lincRNAs that show differing patterns of conservation, including the photo-responsive lincRNA HID1 ((Y. Wang et al. 2014); **Supplemental Figure 7A**), the salt-responsive lincRNA DRIR ((Qin et al. 2017); **Supplemental Figure 7B**), the auxin regulated lincRNA APOLO ((Ariel et al. 2020); **Supplemental Figure 7C**), the pathogen resistance associated lincRNA ELENA (Seo et al. 2017); **Supplemental Figure 7D**), and the cold-responsive lincRNA SVALKA ((Kindgren et al. 2019); **Supplemental Figure 7E**). As expected, a HID1 locus was present in all tested Brassicaceae. Surprisingly, HID1 has an unreported paralog in Arabidopsis, likely resulting from a tandem duplication in the last common ancestor between Arabidopsis and Camelina (**Supplemental Figure 7A**). In addition, multiple syntenic paralogs are retained in species with known whole genome duplication/triplication events (e.g., Brassica and Camelina). The identification of homologs for ELENA and SVALKA were restricted to Lineage I relatives (e.g., Camelina). We identified two unreported APOLO paralogs within the *Arabidopsis thaliana* genome, and multiple paralogs within the *Arabidopsis lyrata* genome, but were unable to find sequence homologs in other Lineage I representatives. None of the *A. lyrata* APOLO homologs were adjacent to the PID1 locus and thus, if expressed, may not be functionally conserved. Finally, we were unable to identify homologs for DRIR1 in any Brassicaceae, suggesting that, based on sequence-level comparisons, it is Arabidopsis-specific.

Some lincRNAs function as transcriptional regulators in cis, influencing the expression of neighboring genes by recruiting Pol-II or transcription factors ((Kopp and Mendell 2018)). For these lincRNAs, conservation of expression next to another gene (collinearity) may be more important than sequence conservation. To address this hypothesis, we used SynMap (Haug-Baltzell et al. 2017) to identify collinear blocks between each of our focal species and then searched for lincRNAs arising from loci adjacent to similar genes within these blocks (see **Materials and Methods**). This approach revealed 1,621 Arabidopsis lincRNAs for which a sequence-dissimilar lincRNA was transcribed in Camelina at a syntenic locus, and an additional 3,560 Arabidopsis lincRNAs that shared synteny with lincRNAs in either Eutrema or Brassica (**Figure 5A**). Interestingly, we identified “transcriptional syntelogs” adjacent to multiple CBF1 loci in Brassica, in a similar orientation and distance to CBF1 as the Arabidopsis lncRNA SVALKA (**Supplemental Figure 7E**). In total, four putative SVALKA transcriptional syntelogs were identified next to CBF1 paralogs in Brassica. A putative transcriptional syntelog was also identified for DRIR1 in Brassica (**Supplemental Figure 7B**). Thus, a significant proportion of “species-specific” lincRNAs in Brassicaceae may in fact be transcribed from syntenic loci with diverged sequence and therefore harbor conserved cis-regulatory functions.

Given the apparent expansion of the HID1 and APOLO gene families, we asked how frequently lincRNA gene families expanded and contracted, particularly in light of whole genome duplication events. We examined lincRNAs for which we were able to identify a sequence homolog in at least one other organism, and then asked if the number of lincRNAs in each gene family matched expectations based on known genomic events (see **Materials and Methods**). For Arabidopsis and Eutrema, which have not undergone recent WGDs, lincRNA gene families are predominantly stable (no change for 89% and 81% of lincRNA families, respectively; **Figure 5B**). In contrast, Camelina and Brassica lincRNA gene families have experienced a significant contraction, with 67% and 98% experiencing a contraction, typically reducing the lincRNA copy number from the expected three copies back to a single copy. This is a higher rate of contraction than has been observed for protein-coding genes (De Smet et al. 2013). Indeed, 71% of Camelina lincRNAs, and 85% of Brassica lincRNAs are single copy, suggesting weak selective pressures to retain these genes in multicopy form. In addition, in Brassica, where the least and most dominant subgenomes have been assigned (Cheng et al., 2013; Tang et al., 2012), most single copy lincRNAs, and most lincRNAs in general, fall within the least fractionated subgenome (LF; n = 26,284), vs the medium fractionated (MF1; n = 21,712) and the most fractionated (MF2; n = 15973; **Supplemental Figure 6F**). For each of these sets of lincRNAs, ~50% are predominantly expressed in datasets examining intra-specific hybrids or RILs (**Supplemental Figure 6F**). These data suggest that lincRNAs, like mRNAs, are preferentially retained on dominant subgenomes following whole genome duplication events, but that lincRNA hybrid-specific expression is not linked to subgenome of origin.

Camelina has a substantial number of multi-copy lincRNA gene families, and thus offers an opportunity to monitor the impact that whole genome duplication events have on lincRNA expression. Camelina is an allohexaploid, consisting of three subgenomes similar to its two progenitor species (*C. hispida* and a *C. neglecta*-like autotetraploid), referred to here as the *C. hispida*, *C. neglecta*, and *C. neglecta* (like) subgenomes. In C. sativa, C. hispida mRNA homeologs are typically more highly expressed relative to those from the other two subgenomes (Chaudhary et al., 2020). To explore how WGD has impacted lincRNA expression, we performed Illumina short read RNA-sequencing in early embryos of Camelina (n = 5). These data were mapped to the reference genome with an updated gene set including our Evolinc-identified lincRNAs. For lincRNAs families with homologs present in the three subgenomes (see **Materials and Methods**), and with at least one member expressed above 1 TPM in Camelina embryos, we see a significant bias against expression of the homologs from the *C. neglecta* (like) subgenome, with similar expression in the other two subgenomes (**Supplemental Figure 8A**). This is in contrast to lincRNAs that are only found in one of the three subgenomes, and thus not evolutionarily related (**Supplemental Figure 8B**). LincRNAs arising from the *C. neglecta* (like) subgenome showed a slight but significant elevated average expression relative to the other two subgenomes (**Supplemental Figure 8B**). Orthologous lincRNAs also behave differently than orthologous protein-coding genes, with *C. neglecta* showing the lowest average expression relative to the other two subgenomes (**Supplemental Figure 8C**). These data suggest that there is an evolutionary context-dependent contrast in expression between lincRNAs found in the same subgenome in Camelina.

We observed a number of distinct features within the Arabidopsis lincRNA dataset, including structured regions, sORFs, and miRNA interaction motifs that may act as functional motifs (Lucero et al., 2020). If these elements are important for lincRNA function, then we would expect them to be conserved. Structural elements, inferred from PIP-seq data, strongly correlated with conservation (**Figure 5C**). Of the 415 lincRNAs for which structured elements were identified, 324 were conserved outside of Arabidopsis. For 70% of these sequence conserved lincRNAs, the structured region was conserved to the same node as the lincRNA itself, suggesting the structural element itself is driving conservation of the lincRNA. An example of this is shown in **Figure 5D**, where the two structural elements of lincRNA “Evolinc_tID.00064432” overlap with deeply conserved (i.e., Node 5) portions of the lincRNA.

We next addressed the degree to which the 158 sORF-containing lncRNAs are conserved, as conservation of the sORF would lend support to the idea that these “lincRNAs” are actually protein-coding transcripts. Of the 127 sORF lincRNAs conserved outside of Arabidopsis, there was no significant variation in the overall rate of conservation relative to non-sORF lncRNAs, indicating that sORF-lincRNAs are not preferentially retained. Of the 158 sORF lincRNAs tested, 53 sORFs were conserved in at least one other species, and 39 were conserved to the same node as the lincRNA from which they were derived (see Materials and Methods; **Figure 5E; Supplemental File 4**). There was no clear bias towards the length of sORF or the encoding transcript (**Supplemental Figure 2**). Some of the conserved sORFs were quite short, such as the sORF within *AT1G06113*, which encodes for a nine amino acid peptide and lies within a region of the sORF-lincRNA that shares almost 100% identity across the 11 species present in the MSA (**Figure 5F**). Although most sORF-lincRNAs (26/36) were previously annotated lincRNAs (i.e., Araport lincRNAs), a subset were identified in this study, suggesting that current filtering schemes are not entirely sufficient for removing short protein-coding transcripts from our dataset.

Finally, we determined the degree to which the predicted miRNA interaction sites within our Arabidopsis lincRNA dataset were conserved. Of the 226 lincRNAs with predicted miRNA interaction sites, 68 were species-specific (**Figure 5G**). A further 83 were sequence-conserved in at least one other Brassicaceae, but the conserved region did not overlap with the putative miRNA interaction site. The remaining 75 lincRNAs contained sequence conserved miRNA interaction motifs, with an example for this shown for AT1G50055 (TAS1B) in **Figure 5H**. Multiple sequence alignments supporting our conservation assignments for structure, sORFs, and miRNA interaction sites can be found in the CyVerse Data Store (**DOI**). LincRNAs with conserved domains are annotated in **Supplemental File 4**. In sum, our evolutionary approach has uncovered conserved lincRNA functional elements that are strong candidates for functional analysis, as well as shed additional light on how plant lincRNAs evolve in the face of WGD.

### Assigning putative function to Brassicaceae lncRNAs

Basic characterization of lincRNA expression, along with an evolutionary analysis, can provide clues as to which lincRNAs in our datasets are potentially functional, but these data alone are not sufficient to begin making functional hypotheses or attributing putative functions. To better clarify when and where the lincRNAs in our catalogs are functioning, we took three approaches. The first was to determine which lincRNAs are stress responsive based on pairwise comparisons of publicly available RNA-seq data (stress vs. control). The second was to use weighted gene co-expression networks (WGCNA) of larger, more complex experiments, to identify modules of similarly expressed protein-coding and lncRNA genes (i.e., guilt-by-association) and infer in which molecular pathway a lincRNA might be acting. Third, as many lincRNAs regulate the expression of neighboring genes (Khyzha et al. 2019; Gil and Ulitsky 2020), we examined correlation of expression of lincRNA-adjacent mRNA gene pairs across tissue and stress expression atlases to identify candidate gene pairs in which the lincRNA might be regulatory.

We first started by annotating which lincRNAs in each species are differentially expressed in response to stress. For Arabidopsis and Brassica, we chose publicly available datasets with multiple independently generated stress experiments (**Supplemental File 4**). In both species, most of the stress-responsive lincRNAs were specific to a particular stress (**Figures 6A, 6B, and Supplemental Figure 9A**), with the highest proportion associated with temperature stress (cold, heat, or cold + heat). We observed a similar pattern for protein-coding genes in both species (**Supplemental Figure 9B and 9C**), therefore we sought to determine which of the Arabidopsis stress-responsive lincRNAs were also ABA responsive. We screened through Arabidopsis RNA-seq data associated with seedlings and roots treated with exogenous ABA (5-100uM) and identified 672 ABA-responsive lincRNAs, 105 of which overlap with our stress-responsive lincRNAs, suggesting these lincRNAs may be stress-responsive in an ABA-dependent manner (**Figure 6A, inset**). As lincRNAs were predominantly responsive to temperature stress (heat and cold), we next asked how many lincRNAs showed an anti-correlated response to temperature stress (i.e., up in heat and down in cold). In both species, heat/cold responsive lincRNAs were predominantly upregulated by heat and repressed by cold (**Figures 6C and 6D**). This pattern was specific for lincRNAs, as most mRNAs were either up in both or down in both conditions (**Supplemental Figure 9D**). Taken together, we observe that a substantial fraction of lincRNAs are differentially regulated during temperature stress in both Arabidopsis and Brassica. Stress and ABA-responsive lincRNAs for each species have been denoted in **Supplemental File 2**. Additionally, all differential expression results from the 4 focal Brassicaceae can be found in **Supplemental File 8**.

WGCNAs help to identify clusters of genes that are coordinated in their expression and thus potentially regulated by, or are regulating, similar pathways. This allows us to assign putative functions to lincRNAs based on significant co-expression with functionally known mRNAs, a process referred to as guilt-by-association (Tian et al. 2008). To remove noise from normalizing across many disparate experiments, we grouped experiments by tissue or, where available, by project (as in the case of the tissue atlases; see Materials and Methods). In total, we identified 987 lincRNAs in Arabidopsis and 3,473 lincRNAs in Brassica whose expression profiles were sufficient to classify them into at least one co-expression module. For example, when we examine the Arabidopsis Klepikova tissue atlas, we identified a module of 233 mRNAs and 11 lincRNAs (8 newly annotated) whose expression peaked in flowers and male reproductive tissues (i.e., anther and pollen; **Figure 6E**). Within this module, gene ontology terms associated with fertilization were enriched, suggesting that lincRNAs within this module are also participating in aspects of fertilization (Module associated GO terms found in **Supplemental Figure 10A**). We also observed lincRNAs falling into co-expression modules within the Klepikova tissue atlas stress experiments. One particular module contains 182 transcripts (176 mRNAs, 6 lincRNAs) whose expression peaks rapidly after wounding (**Figure 6F**). As expected, the GO terms we see with these member mRNAs are highly enriched for response to wounding and jasmonic acid regulation (**Supplemental Figure 10B**), a hallmark hormone response to herbivory and biotic stresses (J. Wang et al. 2020). Thus, through WGCNA and guilt-by-association, we have generated putative annotations for ~1,000 Arabidopsis and ~3,000 Brassica lincRNAs that may guide future *in vivo* functional analyses. These lincRNAs have been annotated with expression modules in **Supplemental File 2**. Detailed WGCNA results can be found in **Supplemental File “All_WGCNA_results”**.

### Expression for a subset of LincRNAs is significantly correlated with adjacent mRNAs

LincRNAs are known to regulate the expression of other genes, either in cis or in trans, through a variety of mechanisms ((Kindgren et al. 2019; Gil and Ulitsky 2020)). One signature of cis-regulatory lincRNAs is correlation in expression relative to neighboring genes across a diverse transcriptomic dataset. To identify putative cis-regulatory lincRNAs, we searched for correlation between all Arabidopsis and Brassica lincRNAs and their immediate neighboring mRNAs that were expressed above 0.1 TPM (i.e., both lincRNA and mRNA > 0.1 TPM) in either their respective tissue atlases or heat stress experiments. In the Arabidopsis tissue atlas we identified 252 lincRNA-mRNA pairs in which both genes in the pair were expressed and for which we could calculate expression correlation (out of 7,875 lincRNAs examined). This correlation was significantly more positive than mRNA-mRNA pairs or random pairs of genes (**Figure 6G**; *P* value=1.62e-14; Wilcoxon rank sum test with Bonferroni multiple testing correction). When examining all genes that fall within 10 Kb of an expressed lincRNA, we observe even stronger positive correlation, in contrast to mRNA-mRNA pairs within the same region, which show very little correlation across all distances measured (up to 10 Kb; **Supplemental Figure 11A**). We observed even more lincRNA-mRNA pairs with correlated expression during heat stress in Arabidopsis (n = 2,544), again with a significant positive correlation relative to mRNA-mRNA pairs (**Figure 6I**). We also observed positive correlation for Brassica lincRNA-mRNA pairs in the Brassica tissue atlas (3,757 out of 23,756 expressed lincRNAs; **Figure 6H and Supplemental Figure 11B**) and heat experiments (n = 6,514), although this correlation was less pronounced than in Arabidopsis. In sum, we identify a subset of lincRNAs whose expression appears to be positively correlated with neighboring genes up to at least 10 Kb away, suggesting that these lincRNAs might be cis-regulatory RNAs. LincRNA-mRNA pairs with a strong correlation (r > 0.5 or r < −0.5), as well as all correlated neighboring pairs are listed in **Supplemental File 2**.

### Synthesizing our functional assignment approach

We searched for experiments that took a holistic approach towards the analysis of stress responsiveness where we could assess the active regulation and response of lincRNAs (i.e., not just RNA-seq, but ChIP-seq or other methods to study the regulation of gene expression). Lee and Serres (2019) performed such an integrative approach to understand hypoxia responses in Arabidopsis seedlings. We set out to re-analyze these data in the context of both mRNAs and lincRNAs. By reanalyzing expression data, we observed 153 Arabidopsis lincRNAs that are differentially expressed in response to hypoxic stress. We also observe 62 lincRNAs that fall into a co-expression module with significant enrichment of GO terms associated with hypoxia (**Supplemental Figure 13A-H**). In Arabidopsis, changes in gene expression in response to hypoxia is regulated predominantly by the transcription factor HRE2 (Hypoxia Responsive Ethylene Responsive Factor 2; AT2G47520). We reexamined Arabidopsis HRE2 ChIP-seq data in the context of the hypoxia stress-associated lincRNAs and found evidence for HRE2 binding to the promoter regions of 20 differentially expressed lincRNAs. We then examined correlation in expression between the HRE2-bound lincRNAs and their adjacent mRNAs, identifying four lincRNAs with positive correlation, one with negative, and two with mixed correlation with adjacent genes (**Figure 6K**). Thus, these seven lincRNAs appear to be specifically regulated by HRE2 in response to hypoxic stress and may act as cis-regulatory elements (annotated in Supplemental File 2).

In sum, we have used a wealth of public data, supplemented with additional short and long-read RNA-seq, to identify and provide putative functional annotations for lincRNAs across four Brassicaceae. We combined our transcriptomics data with comparative genomic and evolutionary analyses to determine conservation of not just the lincRNAs themselves, but also inferred functional elements within the RNAs, such as sORFs, structured regions, and miRNA interaction motifs. Using these approaches, we have identified >100,000 Brassicaceae lincRNAs with multiple lines of functional or contextual evidence that will facilitate downstream functional analyses.

## Discussion

### A comprehensive and unified lincRNA annotation effort for the mustard lineage

Here we generated an expansive catalog of high confidence lincRNAs for four agricultural and model Brassicaceae species by processing > 20,000 publicly available RNA-seq datasets for those species. We supplemented these publicly available data with our own ONT long read sequencing data, and further annotated the identified lincRNAs with epigenetic, genomic, structural, translational, and evolutionary information. These efforts build on previous efforts to catalog novel transcribed elements within plant genomes (Liu et al., 2012; Moghe et al., 2013), and serve as the most exhaustive lincRNA identification and annotation effort to date in any plant species.

Due to the scale of our efforts and the wealth of data available for these four species, we were able to uncover defining features for Brassicaceae lincRNAs, features that may guide future discovery and annotation efforts in other plant lineages. LincRNAs tend to be mono-exonic, but when multi-exonic, harbor longer exons relative to those seen in spliced transcripts. LincRNAs appear to be epigenetically regulated in a distinct manner from both protein-coding genes and transposable elements. And, as expected based on prior observations in plants and mammals, lincRNAs in all four species were, on average, expressed at low levels and displayed significantly higher tissue specificity relative to protein-coding genes in tissue atlases and our ONT data. The exception to this observation were the sORF containing lincRNAs, which behave more similar to protein-coding genes in terms of both higher expression levels and tissue specificity. Interestingly, many of the lincRNAs we identified displayed high expression in, or were restricted to, very specific cell types (e.g., meristematic tissue) or experimental conditions (e.g., environmental stress) suggesting that 1) lincRNA expression is highly context and cell-type specific, and 2) sampling bulk tissues may not accurately reflect a lincRNA’s contribution to the transcriptome. The lincRNAs restricted to inter-accession crosses as in *B. rapa* may be the result of improper transcriptional control given their relatively even distribution across the genome or, albeit less likely, may reflect transcripts that help mediate compatibility of two subtly different genomes.

### Using comparative genomics to provide functional insights

Given that we identified thousands of lincRNAs in each of our four focal species, functional analyses will need to be prioritized. In order to facilitate that prioritization, we used a comparative genomic approach to assess the degree to which each identified lincRNA is conserved, and if there are any particular motifs of interest within those conserved lincRNAs. As expected based on prior observations in plants and mammals, we observed low levels of sequence conservation for lincRNAs identified in each of the four species relative to protein-coding genes. However, when sequence homologs were detected between two species (e.g., Arabidopsis to Brassica), those sequence homologs were predominantly annotated as lincRNAs and not protein-coding genes. Inspired by a smaller comparison between Arabidopsis and Aethionema (Mohammadin et al., 2015), we also searched for and observed a cohort of lincRNAs that are transcriptional syntelogs in that they are transcribed from similar genomic positions in multiple species but share little sequence conservation. LincRNAs that regulate gene expression in cis are an interesting class of transcripts from an evolutionary perspective in that positional and transcriptional conservation may be more critical than sequence conservation. Although additional study is needed, we posit that these lincRNAs may be functionally conserved in regulating expression of the orthologous genes to which they are adjacent in each species. An exciting set of candidates for further study are the putative SVALKA transcriptional syntelogs we identified in Brassica. In Arabidopsis, SVALKA regulates an adjacent, non-overlapping, protein-coding gene through transcriptional interference. This mode of function in particular might depend more on conservation of transcription, and from where transcription arises, than it does on sequence similarity.

Identifying lincRNAs in species with recent WGD events (e.g., Camelina and Brassica) allowed us to more closely examine how lincRNAs evolve following these genomic events. LincRNAs are not typically retained as multicopy loci following WGD events. In Brassica, lincRNAs are predominantly retained as single copy from the least fractionated genome. The fractionation - and retention, of a certain set of lincRNAs may suggest functional interactions (e.g., genetic or molecular) are preferentially retained following WGD events - similar to that observed for protein-coding genes (Emery et al., 2018; Schnable et al., 2012). When lincRNAs are retained as multicopy, their expression appears to be more sensitive to the influence of subgenome dominance than protein-coding genes - perhaps explaining why they are fractionated from the genome. However, the retention, and expression, of paralogous lncRNAs such as *HID1* may suggest that lncRNAs can be functionally retained post-duplication in a similar manner as protein-coding genes. Further studies are needed to determine if these paralogous lincRNAs (e.g. *HID1*) have sub or neo-functionalized as is often the case for retained proteins.

Using multiple sequence alignments for our sets of conserved lincRNAs, we also determined if the identified structural, putative miRNA binding, or sORFs were within those conserved regions. Although we did identify examples of conserved sORFs, to our surprise we did not observe strong correlation between sORF and lincRNA conservation. One particularly interesting conserved sORF is found within the lincRNA AT1G06113. The Ribo-seq identified sORF within this lincRNA is only nine amino acids long, but is almost perfectly conserved across the Brassicaceae and even in *T. hassleriana*. The functional significance of this peptide, as well as the other lincRNA-derived small proteins remains to be determined. In contrast to the sORF-containing lincRNAs, the regions we identified to be protein-bound and structured were typically conserved to the same degree as the lincRNA itself. This conservation suggests these structured regions are important for function and may bind similar proteins in multiple species. Thus, identifying the protein binding partner in Arabidopsis might help provide functional insights for these lincRNAs across the family as well as develop a protein-RNA interaction database for improving functional predictions.

### Using omics-approaches to assign putative function to Brassicaceae lincRNAs

Our ultimate goal, beyond identifying lincRNAs in each of these species, was to annotate these lincRNAs so as to aid in future functional studies. We used expression data to assign lincRNAs into broad regulatory categories, such as stress-responsive, cis-regulatory, or others associated with GO-terms extracted from network analyses. As most functionally described lincRNAs to date are associated with changes in the environment (i.e., biotic/abiotic stress), our initial expectations were that most lincRNAs would be stress responsive. Interestingly, this was not the case. Roughly 10% of the lincRNAs identified in Arabidopsis and Brassica are stress-responsive, with most responding to temperature stress. While this could be linked to changes in genome-wide epigenetic control that is not specific to lincRNAs, there does appear to be a degree of response specificity. A majority of the temperature (cold or heat) responsive lincRNAs were either specific to one stress or the other, or showed opposite responses to the two stresses. Furthermore, we also identified a set of lincRNAs whose response appears to be ABA-dependent. The preponderance of lincRNAs associated with temperature stress in our dataset may simply reflect sampling bias as our analyses were dependent on publicly available data. However, given the lincRNAs and NAT-lncRNAs that have already been functionally described as temperature responsive in Arabidopsis (Kindgren et al., 2018; Castaings et al., 2014; Zhao et al., 2018), the potential for widespread adaptation to environmental conditions by lincRNAs remains an exciting avenue for future research.

## Methods

### Plant materials and growth

*Arabidopsis thaliana* (Col-0; (Lamesch et al. 2012), *Brassica rapa* (R-0-18; (Howe et al. 2021), *Camelina sativa* (cultivar Ames), and *Eutrema salsugineum* (Shandong; (Yang et al. 2013) seeds were surface sterilized by washing with 70% ethanol followed by soaking in 30% bleach and 1% Tween 20 for 10 minutes before being rinsed and plated on ½ MS media supplemented with 0.5% sucrose. Plates were placed in the dark at 4°C for 5 days before being moved to a long day (16 hour light 22°C/8 hour dark 20°C) growth chamber. Ten days after germination, seedlings were either collected in liquid nitrogen or transplanted to soil and placed into the same growth chamber. For leaf samples, leaves were either collected 4 weeks after germination, or at the mature most vegetative stage, whichever came first. Finally, for flower samples, opened flowers with no sign of developing fruit were collected. All plant samples were immediately frozen in liquid nitrogen and stored in a −80°C freezer until ready for processing.

### RNA extraction and ONT library preparation

Frozen plant samples were pulverized in liquid nitrogen using a chilled mortar and pestle until a fine powder was obtained. RNA was extracted using the RNeasy Plant Mini kit (Qiagen) following the manufacturer’s instructions. Purified RNA was used as input for the Dynabeads mRNA Purification kit (Invitrogen). Purified poly-A RNA was used as input for the Nanopore direct cDNA sequencing kit (SQK-DCS109) following the manufacturer’s instructions. Nanopore libraries were sequenced on a MINion sequencer (R9.4.1 flowcell). Raw reads were basecalled using a GPU-enabled version of Guppy in the command line.

### Illumina RNA-sequencing of Camelina sativa seeds

Developing seeds of four *Camelina sativa* accessions were collected in biological triplicate at ~15 days post anthesis and immediately placed in liquid nitrogen. Total RNA was isolated from developing seeds using the PureLink® Plant RNA Reagent (Thermo Fisher Scientific, Waltham, MA, USA) and its associated protocol. Extracted RNA was then purified further using an RNeasy RNA clean-up kit (Qiagen, Valencia, CA, USA) and quantified on a Qubit fluorometer (Life Technologies, Carlsbad, CA, USA). Sequencing libraries were prepared with the SENSE mRNA-seq library prep kit and protocol, using up to 1,000 ng total RNA per sample (Lexogen GmbH, Vienna, Austria). Individual transcriptome libraries were quantified using a Qubit fluorometer and fragment size, distribution, and overall library quality was determined with an Agilent Bioanalyzer (Agilent, Santa Clara, CA, USA) system. Samples were pooled into three final libraries and sequenced by Novogene (Sacramento, CA, USA) on an Illumina HiSeq platform (Illumina, San Diego, CA, USA) producing 150 bp paired-end reads.

### LincRNA identification and basic characterization

The RMTA (Peri et al. 2020) pipeline was used to process all available short read RNA-seq experiments as of December 2018 within the CyVerse Discovery Environment (Merchant et al. 2016) using the HiSat2 and Stringtie (Pertea et al., 2016) mapping and assembly options. Assembled transcripts were then processed through the Evolinc (Nelson et al. 2017) pipeline to identify lincRNAs. For *Arabidopsis thaliana*, the TAIR-10 assembly was used as a reference for the initial RMTA workflow (including mapping, quantification, and transcript assembly), for *Brassica rapa* the Ensembl v1.0, for *Camelina sativa* v2.0 from Ensembl (Plant Release 51), and for *Eutrema salsugineum* Phytozome v1.0 (Yang et al. 2013). An updated annotation including newly identified lincRNAs for each species can be downloaded from the CyVerse Data Store: (**DOI from CyVerse**).

Basecalled Nanopore reads were demultiplexed and processed following (A Eccles 2019). To identify lincRNAs with Evolinc, processed reads were aligned to each species’ genome using Minimap2 (H. Li 2018). Mapped reads were assembled into transcripts using Stringtie2 (Kovaka et al. 2019) using the -L parameter. Transcript assemblies were then used as input for Evolinc for lincRNA identification.

The BEDTools suite (Quinlan and Hall 2010), “nuc” function) was used to characterize the GC content and gene lengths of mRNAs and lincRNAs. Exon counts were determined using the R (Team and Others 2013) R Core Team, 2013, version 4.1.0) package GenomicFeatures (Lawrence et al. 2013), “exonsBy” function, v 1.44.1).

### Analysis of DNA methylation patterns and histone modification dynamics

LincRNA and mRNA epigenetic profiles were monitored by reprocessing publicly available whole genome bisulfite sequencing (WGBS) datasets as well as chromatin immunoprecipitation with sequencing (ChIP-seq) experiments (see **Supplemental File 6**). WGBS data was processed using the Bismark tool (Krueger and Andrews 2011) with default parameters to generate BedGraph files which were then converted to bigWig format. These bigWig files were used in deepTools (Ramírez et al. 2014) with the “computeMatrix’’ function and “scale-regions’’ option to visualize CpG methylation over lincRNA and mRNA genes. ChIP-seq datasets were processed by first aligning the raw reads to the respective genome using BWA-MEM (H. Li 2013) which is deployed as a CyVerse app in the Discovery Environment (BWA_mem_0.7.15 with default settings). SAM files from BWA-MEM were converted to sorted BAM files. The Picard Toolkit (Picard Toolkit 2019) was used to remove PCR duplicates using the “lenient” setting for the “VALIDATION_STRINGENCY” option. These processed BAM files were then used as input for the deepTools “bamCompare’’ function with a ChIP input sample (if available) as a comparison experiment. For Arabidopsis, paired RNA-seq experiments were used to determine which genes were expressed for plotting for WGBS and ChIP-seq experiments. For Brassica, the defined tissue atlas was used instead.

### Characterization of lincRNA expression patterns

To characterize expression from ONT-seq data, Minimap2 was used to map ONT-reads to transcriptomes for each of the respective species’ updated gene sets (prior annotated genes + Evolinc lincRNAs) using similar parameters as above. Minimap2 produced BAM files were used as input for Salmon in alignment-based mode, specifying the --noErrorModel option (Patro et al. 2017)Soneson et al. 2019, (Patro et al. 2017). TPM values were aggregated from each experiment using the tximport R package (Soneson, Love, and Robinson 2015) to obtain gene level expression estimates.

Specific Illumina short read datasets from Arabidopsis and Brassica were used to gain additional resolution of tissue specific expression. For Arabidopsis, the Klepikova *et al.,* 2016 (NCBI PRJNA314076) tissue expression atlas was reprocessed, and for Brassica two datasets were combined to create a tissue atlas similar to Arabidopsis (PRJNA253868 & PRJNA185152). RNA-sequencing reads (FASTQ) associated with each dataset were re-aligned to transcripts with Salmon using XXX parameters to generate transcript-level expression values (TPM). Gene level expression values were obtained as above using tximport. To calculate the tissue specificity metric τ TAU, TPM values were first averaged across replicates. TAU was then calculated as described by (Yanai et al. 2005) using quantile normalized TPM values generated from the preprocessCore R package (Bolstad n.d.). To assess tissue of maximum expression, variance stabilized transformed expression values generated from DESeq2 were utilized (Love, Huber, and Anders 2014).

The DESeq2 package (in R), with the DESeq and results functions, were used to identify differentially expressed genes in pair-wise comparisons. For time-course studies, only the first and last treatments were examined, treating each of them as separate analyses (e.g. early stress response vs. late stress response). Genes were considered to be differentially expressed if they had a log_2_ fold change (L_2_FC) greater or less than 1 or −1, respectively, as well as an adjusted p-value (q-value, FDR) of 0.05 or lower.

### Analysis of co-expression modules

To assess co-expression modules, raw RNA-seq data from select stress experiments (**Supplemental File 6**) was re-processed using Salmon as performed above. The tximport and DESeq2 R packages were used to import Salmon quantification files and convert expression estimates to a variety of normalized values (TPM, VST, normalized counts, etc.) The R package CEMiTool (Russo et al. 2018), v 1.10.2) was used to perform gene co-expression network analyses. In most cases, log_2_ + 1 converted TPM values obtained from tximport were used as input to CEMiTool. Gene ontology (GO) information was provided to CEMiTool from the biomaRt R package (Durinck et al. 2005), and protein-protein interaction data was obtained from the Arabidopsis Protein Interaction Network (Brandão, Dantas, and Silva-Filho 2009).

### Measuring expression correlation of adjacent genes

Arabidopsis and Brassica expression data from the above described tissue atlases or heat experiments (Arabidopsis: PRJNA324514, Brassica: PRJNA298459) were normalized using the DESeq2 “vst” function with the “blind” parameter set to false. Additionally, for the tissue atlases, replicates for each tissue were averaged, if applicable. Genes that did not vary substantially across the input experiments were removed. This was performed by calculating the interquartile range for expression of all genes and only those in the top 50% of IQR values were retained (50% most variable genes). Pearson correlation coefficients of expression were then calculated between all remaining genes post filtering using the corrr R package (Kuhn and Wickham 2020), v 0.4.3). Relevant correlations were then filtered for between lincRNAs and their nearest upstream and/or downstream mRNA neighbors. LincRNA-mRNA pairs separated by fewer than 100 base pairs were removed before subsequent analyses. Random gene pairs were generated from all pairwise correlations using the slice_sample function from the dplyr R package.

To analyze all gene pairs within defined distances, the bedmap function from the BEDOPS suite (Neph et al. 2012) was used with the range, echo, and echo-map-id options. This generated all lincRNA-mRNA or mRNA-mRNA pairs within 200, 500, 1000, 2000, 5000, and 10000 base pairs of each other. These gene pairs were used to filter out the pairwise correlations generated above.

### Assessing lincRNA function from multi-omics hypoxia datasets

The multi-omics datasets generated by Lee and Bailey-Serres (2019) were re-processed using the above methods. After processing ChIP-seq data as above, the output BAM files were used as input for HOMER motif analysis (Heinz et al. 2010) following along closely with the provided “Next-Generation Sequencing Analysis” tutorial provided by HOMER. Differentially expressed genes were generated using a basic design strategy in DESeq2. Briefly, the early hypoxia stress was compared to the early control samples (2 hour hypoxia vs 2 hour control using the contrasts option with the results command from DESeq2), and the later hypoxia stress was compared to the later control experiments. Differentially expressed genes for the reoxygenation experiments were not analyzed, but the expression data was used for constructing the DESeq data set and running the differential expression analysis. For identifying relevant co-expression modules including lincRNAs that may be involved in the hypoxia response, CEMiTool was used as above using log transformed normalized counts from DESeq2.

### Identifying translated sORFs from Ribo-seq

Translated sORFs within the lincRNAs were identified using our recent Ribo-seq and RNA-seq data in Arabidopsis seedling (GEO accession no. GSE183264; (Wu and Hsu 2021). Briefly, BAM files of the Ribo-seq and RNA-seq and a GTF containing the lincRNAs and Araport11 annotated genes were imported into RiboTaper (Calviello et al., 2015). The Ribo-seq read lengths and offsets for RiboTaper were 24, 25, 26, 27, 28 and 8, 9, 10, 11, 12, respectively, as previously described (Wu and Hsu 2021). RiboTaper then computed 3-nucleotide periodicity, which corresponds to translating ribosomes move 3-nucleotide per codon, in each possible ORF within the transcripts. The sORFs were considered translated if they displayed significant 3-nucleotide periodicity and the translated ones were extracted from the RiboTaper output ORF_max_filt file.

To identify lincRNAs harboring putative sORFs based on mass spectronomy data, proteomic experiments, PXD026713 and PXD009714, were retrieved from the PRIDE repository. Raw chromatograms were analyzed using MaxQuant software (Version 1.6.0.16) with Andromeda - an integrated peptide search engine (Cox at al., 2011). Following search settings were applied: a maximum of two missed cleavages was allowed, and the threshold for peptide validation was set to 0.01 using a decoy database. In addition, methionine oxidation and N-terminal acetylation were considered variable modifications, while cysteine carbamidomethylation was a fixed modification. The minimum length of a peptide was set to at least seven amino acids. Moreover, label-free protein quantification (LFQ) was applied. Peptides were identified using the Araport 11 database (The Arabidopsis Information Resource, www.Arabidopsis.org) and a library of all Arabidopsis lincRNA ORFs (positive strand) obtained using Transdecoder.

### Evolutionary analyses

LincRNA sequence homologs were identified using the Evolinc-II module (v2.0, https://github.com/Evolinc/Evolinc-II; e-value of −10), with the following genomes: *Arabidopsis thaliana* (TAIR10), *Arabidopsis lyrata* (Ensembl v1.0, Hu et al., 2011), *Capsella grandiflora* (Phytozome v1.1, Slotte et al., 2013), *Capsella rubella* (Phytozome v1.1, Slotte et al., 2013), *Camelina sativa* (Ensembl v2.0), *Cardamine hirsuta* (v1.0, Gan et al., 2016), *Brassica rapa* (Ensembl v1, Wang et al., 2011), *Schrenkiella parvula* (Phytozome v2.0, Dassanayake et al., 2011 and Oh et al., 2014), *Eutrema salsugineum* (Phytozome v1.0, Yang et al., 2013), *Aethionema arabicum* (CoGe vVEGI 2.5. gID 20243, Haudry et al., 2013 and Nguyen et al., 2019), *Tarenaya hassleriana* (CoGe v4, gID 20317, Cheng et al., 2013). For each of the four species, the entire lincRNA list (LC + HC) were included as query in the analyses. LincRNAs were determined to be restricted to a particular node if no sequence homolog was identified in a more distantly related species. LincRNAs were determined to be conserved as lincRNAs or mRNAs in other species if they overlapped by 50% or more with an annotated gene on the same strand. If not, they were considered to be unannotated. Multiple sequence alignments produced by Evolinc-II (using MAFFT) were imported into Geneious (Genious Prime 2021.1.1, https://www.geneious.com) for downstream structure, sORF, and miRNA motif analysis.

Transcriptional syntelogs were identified by downloading the DAGChainer output, with genomic coordinates, from pairwise SynMap analyses between Arabidopsis and each of the three other species (links to regenerate analyses: Camelina https://genomevolution.org/r/1fjg7, Eutrema https://genomevolution.org/r/1f7si, and Brassica https://genomevolution.org/r/1f79g). LincRNAs that were found within syntenic blocks (10 colinear protein-coding genes), between orthologous genes in either of the pairwise SynMap analyses, and in the same orientation to at least one of the neighboring orthologous genes were considered transcriptional syntelogs. To infer lincRNA gene family contraction or expansion, a rudimentary ancestral state reconstruction was performed. For Arabidopsis, ancestral gene copy number for each Arabidopsis lincRNA was inferred by averaging the number of recovered sequence homologs in (at minimum) *A. lyrata*, *C. rubella*, and *C. grandiflora*. Species-specific lincRNAs were not examined. For Camelina, *C. rubella*, *C. grandiflora*, *A. thaliana*, and *A. lyrata* were used to determine the copy number in the last common ancestor. This value was then multiplied by three (to account for the Camelina-specific whole genome triplication event). Values above or below this value were considered to be expansions or contractions, respectively. A similar approach was performed for Brassica and Eutrema.

MSAs were manually scanned to infer depth of conservation of sORFs, putative miRNA binding motifs, and structural/protein-binding elements. On top of lincRNA sequence homology and synteny requirements, for a sORF to be considered conserved, the start and stop sites within the annotated Arabidopsis lincRNA must be positionally conserved (within +/- three AA). In addition, the translated amino acid sequence must be 75% identical in pairwise alignments between Arabidopsis and each putative homologous sORF. To identify putative miRNA binding sites, all lincRNAs were scanned for motifs using psRNATarget (Dai et al., 2018) using an expectation score of 2.5 as cutoff. LincRNAs with putative miRNA binding motifs were then compared against the list of lincRNAs that were conserved outside of Arabidopsis. MSAs were then scanned for the presence of miRNA motifs. Motifs with complete coverage and no more than two (pairwise) mismatches in at least one other species were considered for evolutionary comparisons. For conservation of structural/protein-binding motifs, structured regions inferred by PIP-seq (GEOs GSE58974 and GSE86459; Gosai et al., 2015 and Foley et al., 2017) were intersected with lincRNAs using Bedtools intersect (Quinlan et al., 2010). Arabidopsis lincRNAs, their sequence homologs (from Evolinc-II) and structured regions were combined into a MSA using MAFFT (Nakamura et al., 2018) for manual inspection. PIP-seq motifs were considered conserved if the entire motif was contained within an alignable region of a sequence homolog from another species. For a motif (sORF, miRNA, or structural) to be considered conserved to a particular node, at least one species that shares that node with Arabidopsis was required to contain those motifs under the parameters described above.

## Author Contributions and Acknowledgements

KRP, EL, MAB, and ADLN developed the project. KRP, LY, LW, ES, and ADLN performed the analyses. ACND performed RNA extractions and ONT-sequencing. JRB performed the Camelina RNA-seq. LW and PH performed the Ribo-sequencing. ES and AS examined the MS data. KRP, EL, MAB, and ADLN wrote the manuscript. The authors would like to acknowledge the NSF Graduate Research Fellowship Grant DGE-1746060 (awarded to K.R.P.), NSF-IOS grant 1758532 (awarded to ADLN), NSF-IOS 1444490 (awarded to EL and MAB), and NSF-IOS 2023310 (awarded to ADLN and EL).

## Supporting information

Supplemental File 1: List of SRAs examined for all four species

Supplemental File 2: Functional annotations for each lincRNA from all four species

Supplemental File 3: Evolinc II results for all four species

Supplemental File 4: sORF and structural motif conservation and characteristics

Supplemental File 5: Predicted miRNA binding motifs for all four species

Supplemental File 6: SRAs, with associated metadata, used in targeted transcriptomic studies

Supplemental File 7: Araport11 lincRNAs that were removed from analysis

Supplemental File 8: Expression information for mRNAs and lincRNAs in paired analyses

## Figure Legends

**Supplemental Figure 1:**
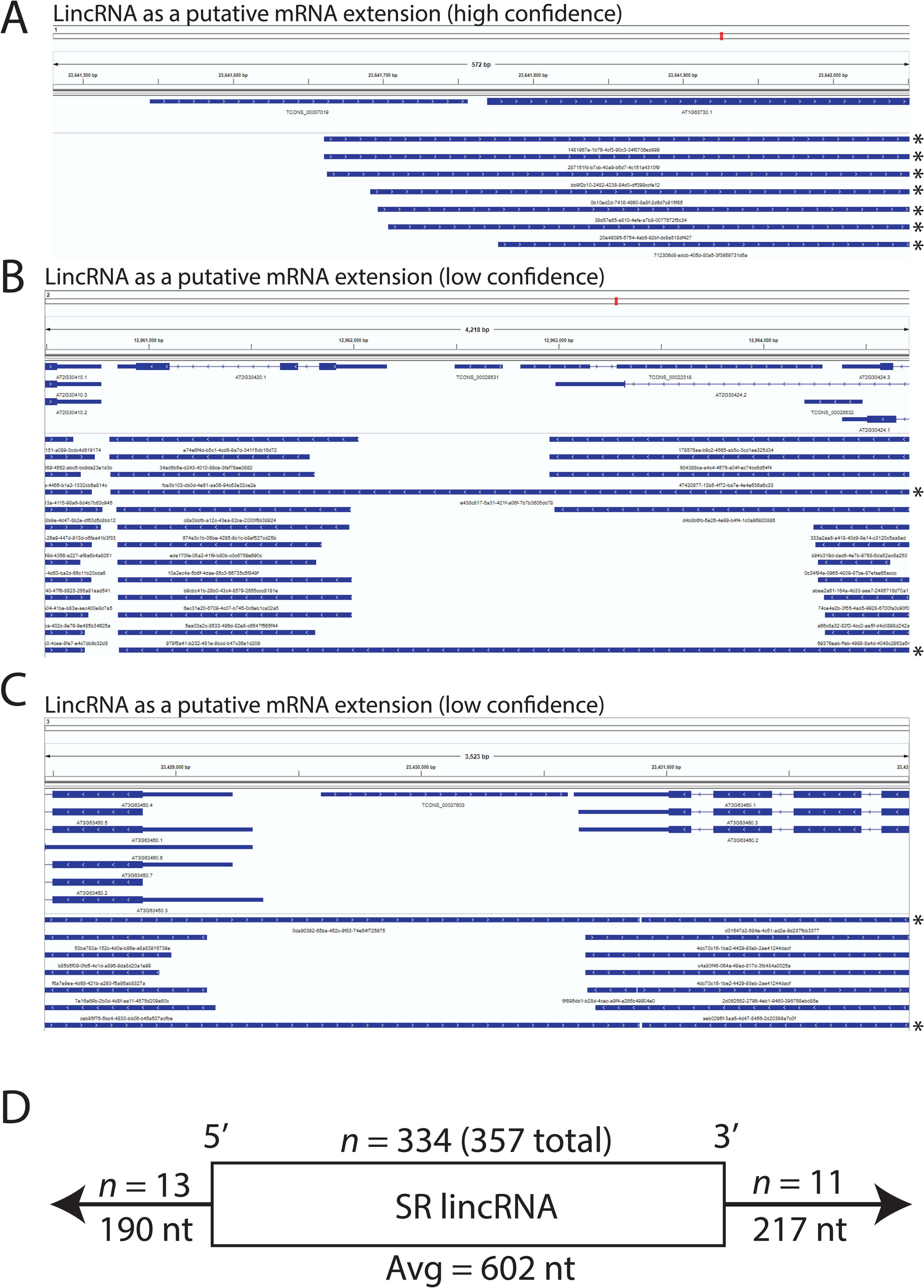
Assessing the assembly quality of Arabidopsis lincRNAs. **A**) Illumina short read RNA-seq lincRNA which was reassessed as an UTR extension of a neighboring mRNA based on Nanopore long read sequencing. **B-C**) Two different lincRNAs initially believed to be mRNA associated, but upon closer inspection were miscalled due to apparent genomic DNA contamination in the ONT-sequencing data. **D**) Comparing the annotated gene structure of lincRNAs assembled in both long and short sequencing reads (n = 357). 334 of the lincRNAs assembled in both technologies were in complete agreement regarding 5’ and 3’ positions, as well as exon structure. 13 lincRNAs were annotated as being, on average, 190 nt longer by ONT-sequencing, whereas 11 were annotated as being 217 nt longer in the 3’ direction.

**Supplemental Figure 2:**
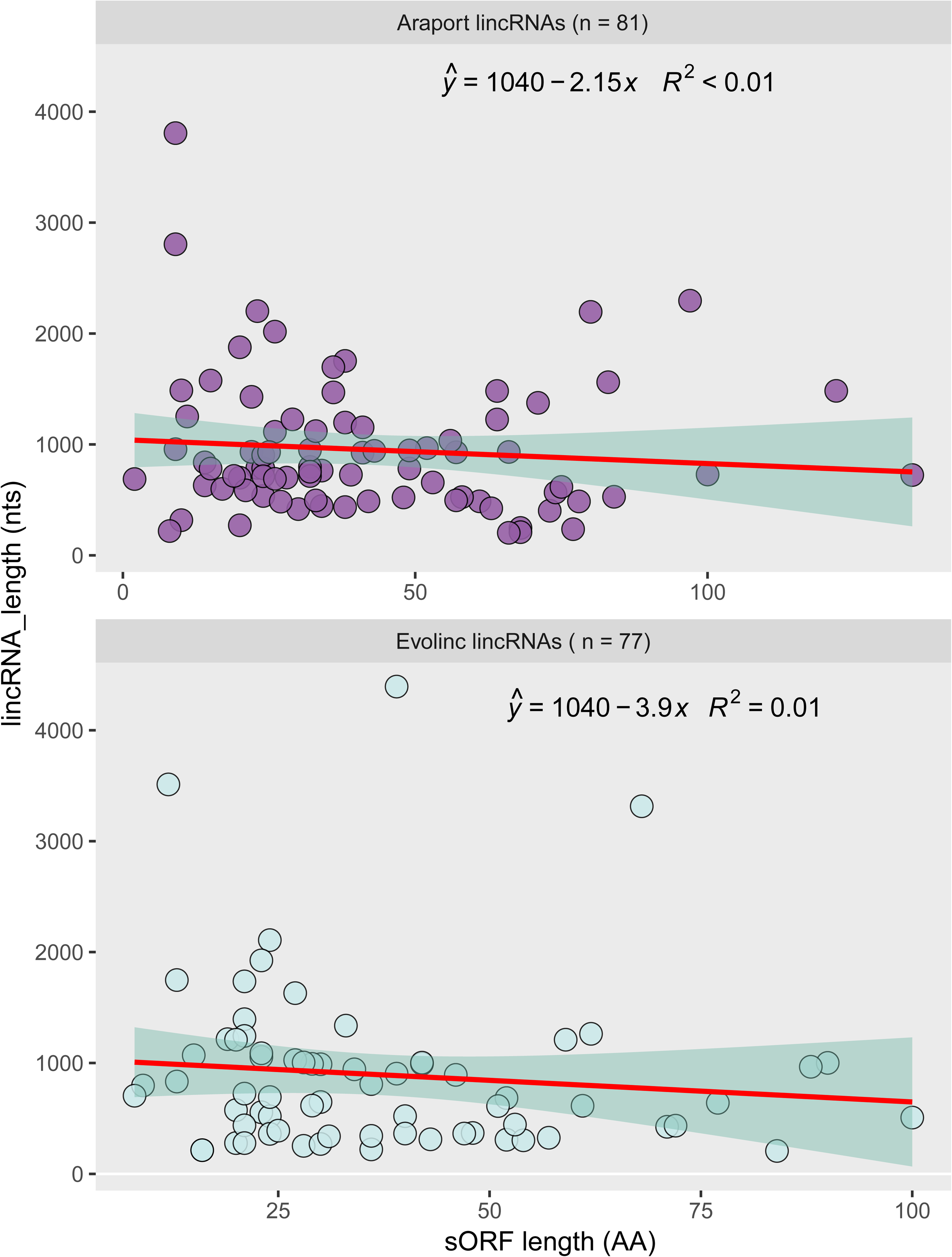
Scatterplots describing the lack of correlation between sORF length and lincRNA transcript length for Araport (top) lincRNAs and Evolinc (bottom) lincRNAs.

**Supplemental Figure 3:**
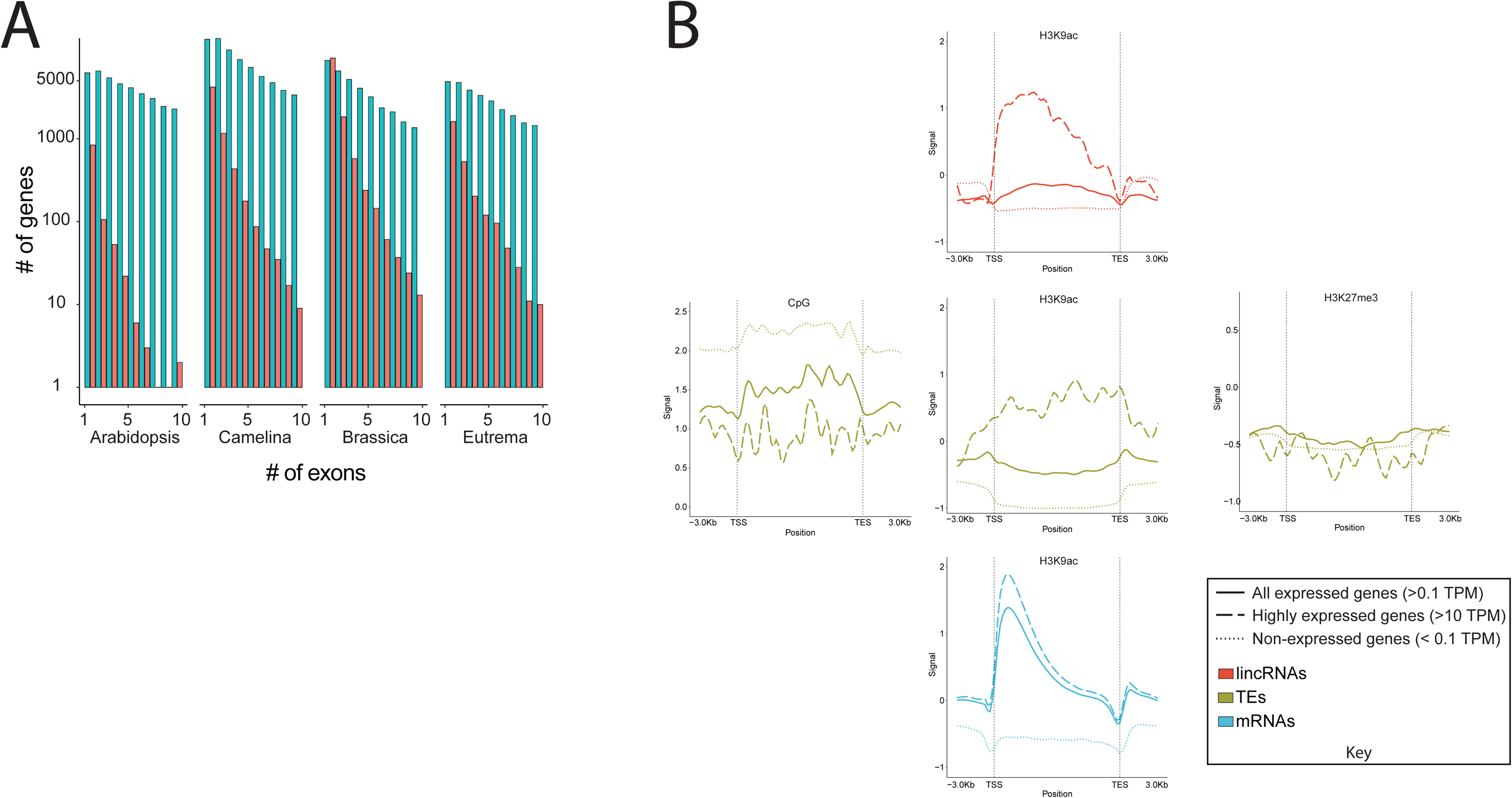
Further basic characterization of lincRNAs. **A**) Exon per transcript distribution of lincRNAs and mRNAs in each of the 4 focal species (mRNAs in blue and lincRNAs in red.) **B**) Metagene plots of CpG and H3K27me3 for transposable elements, as well as H3K9 acetylation for lincRNAs, mRNAs, and TEs. Transcripts are separated based on expression from paired RNA-seq data.

**Supplemental Figure 4:**
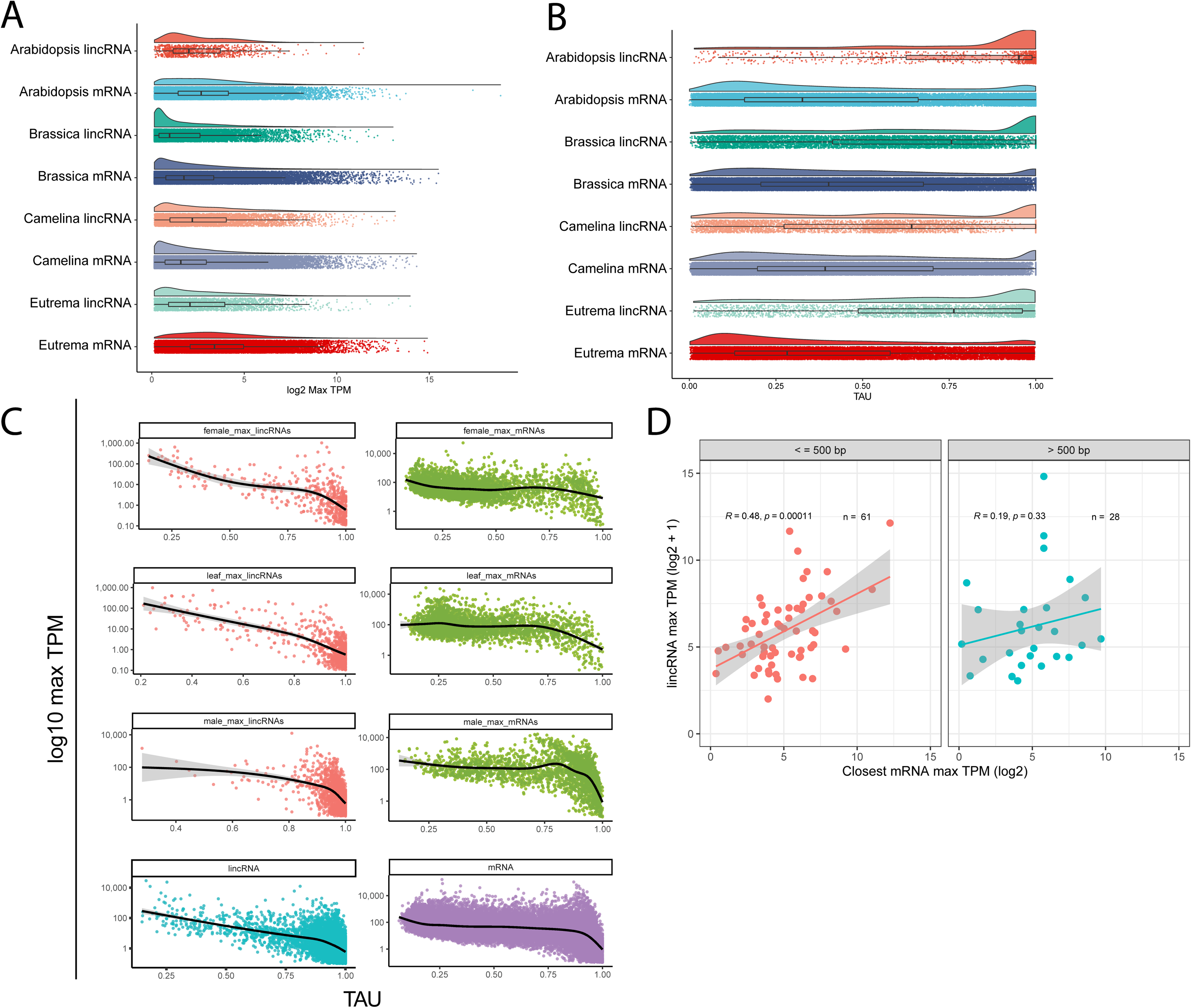
Additional expression characteristics. **A**) ONT-sequencing derived maximum TPM values for lincRNAs and mRNAs for each focal species. Each species’ mRNA-lincRNA comparison is significantly different at *P* < 1.0e-9 using a pairwise Wilcoxon rank sum test with Bonferroni multiple testing correction. **B**) Tissue specificity comparisons from Nanopore TPM values for mRNAs and lincRNAs from all four species. Each species’ mRNA-lincRNA comparison is significantly different at *P* < 2e-16 using a pairwise Wilcoxon rank sum test with Bonferroni multiple testing correction. **C**) Relationship between maximum expression (TPM) and tissue specificity (TAU) between all expressed Arabidopsis lincRNAs (left) and mRNAs (right) within female reproductive tissues (top) leaf tissue, male reproductive tissue, and all tissues combined (bottom). Black lines represent the best fit line of the data. **D**) Relationship between expression levels of low TAU (broadly expressed) lincRNAs and their neighboring mRNAs divided into two groupings based on distance to closest mRNA.

**Supplemental Figure 5:**
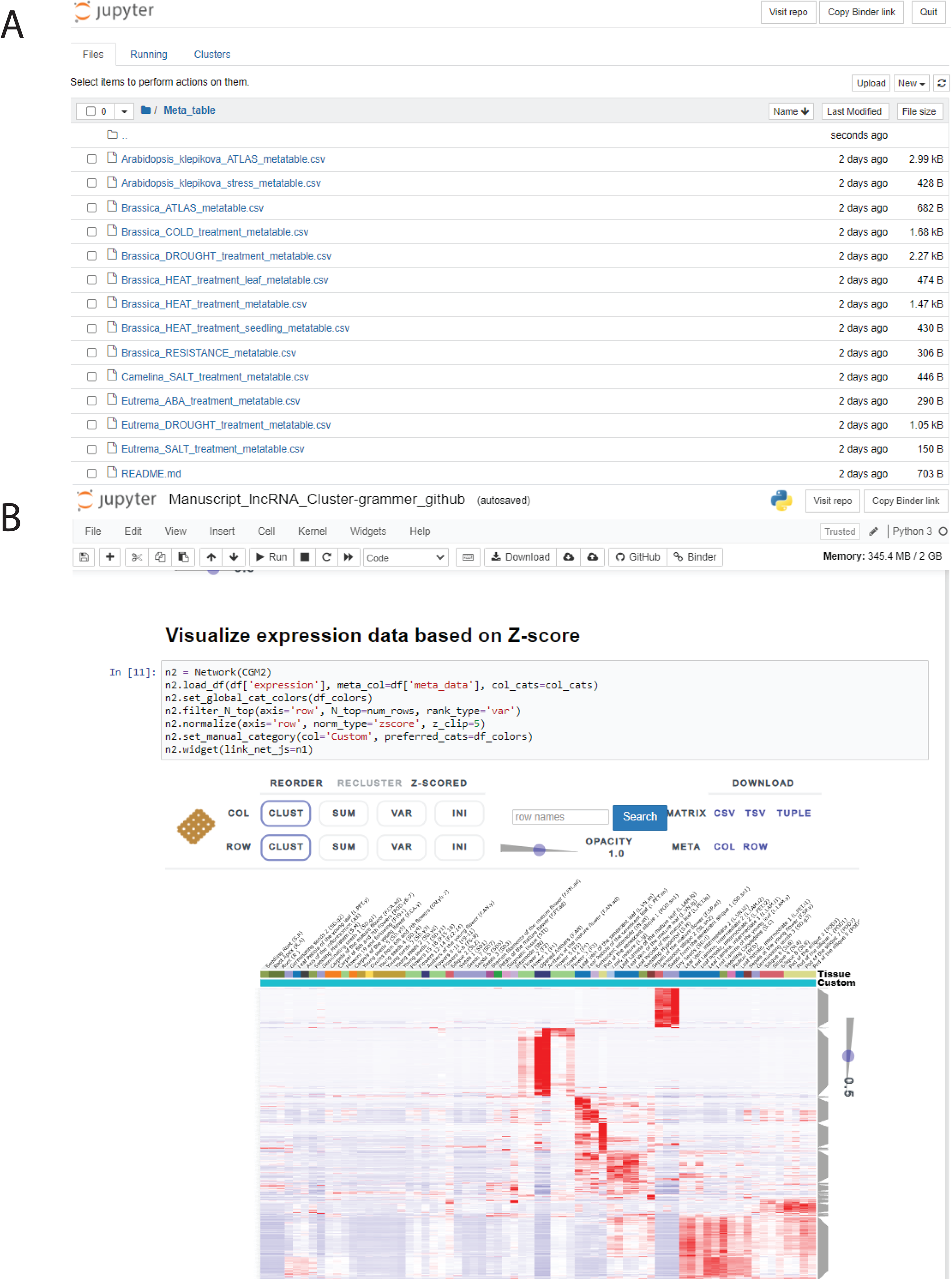
Example screenshot of Clustergrammer Jupyter notebook in which users can examine normalized expression values for mRNAs and lincRNAs across multiple stress and tissue atlases. See https://github.com/Evolinc/Brassicaceae_lincRNAs.

**Supplemental Figure 6:**
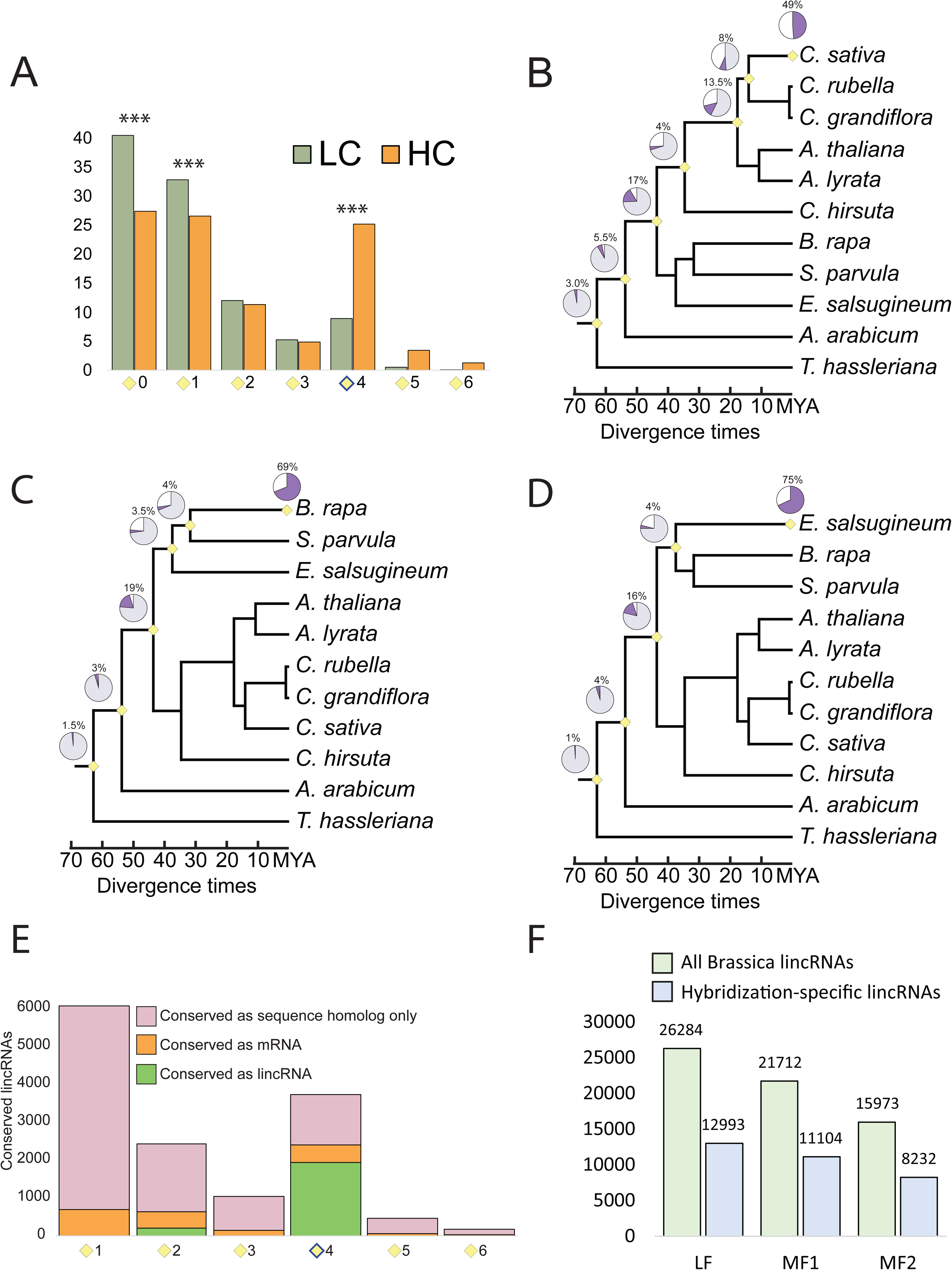
Evolutionary features of Brassicaceae lincRNAs. **A**) Percent of low-confidence and high-confidence Arabidopsis lincRNAs that are sequence conserved at each evolutionary node. Asterisks denote significant difference between observed homolog recovery for the two classes of lincRNAs (p-value <<< 0.01; Student’s t-test). **B-D**) Conservation of lincRNAs from Camelina (**B**), Brassica (**C**), and Eutrema (**D**) across representative Brassicales. The purple wedge in the pie chart in each panel represents the percent of lincRNAs for which sequence homologs were recovered at each node, thus indicating that each lincRNA was conserved to at least that node. **E**) Number of Arabidopsis lincRNAs for which the sequence homolog corresponded to another lincRNA (green bar), mRNA (orange bar) or unannotated sequence (pink bar) at each particular node. **F**) Number of lincRNAs originating from each of the Brassica rapa subgenomes (LF = least fractionated, MF1 = medium fractionated, and MF2 = most fractionated). Coordinates for determining location of subgenomes within the Brassica genome were obtained from Cheng et al., 2013.

**Supplemental Figure 7:**
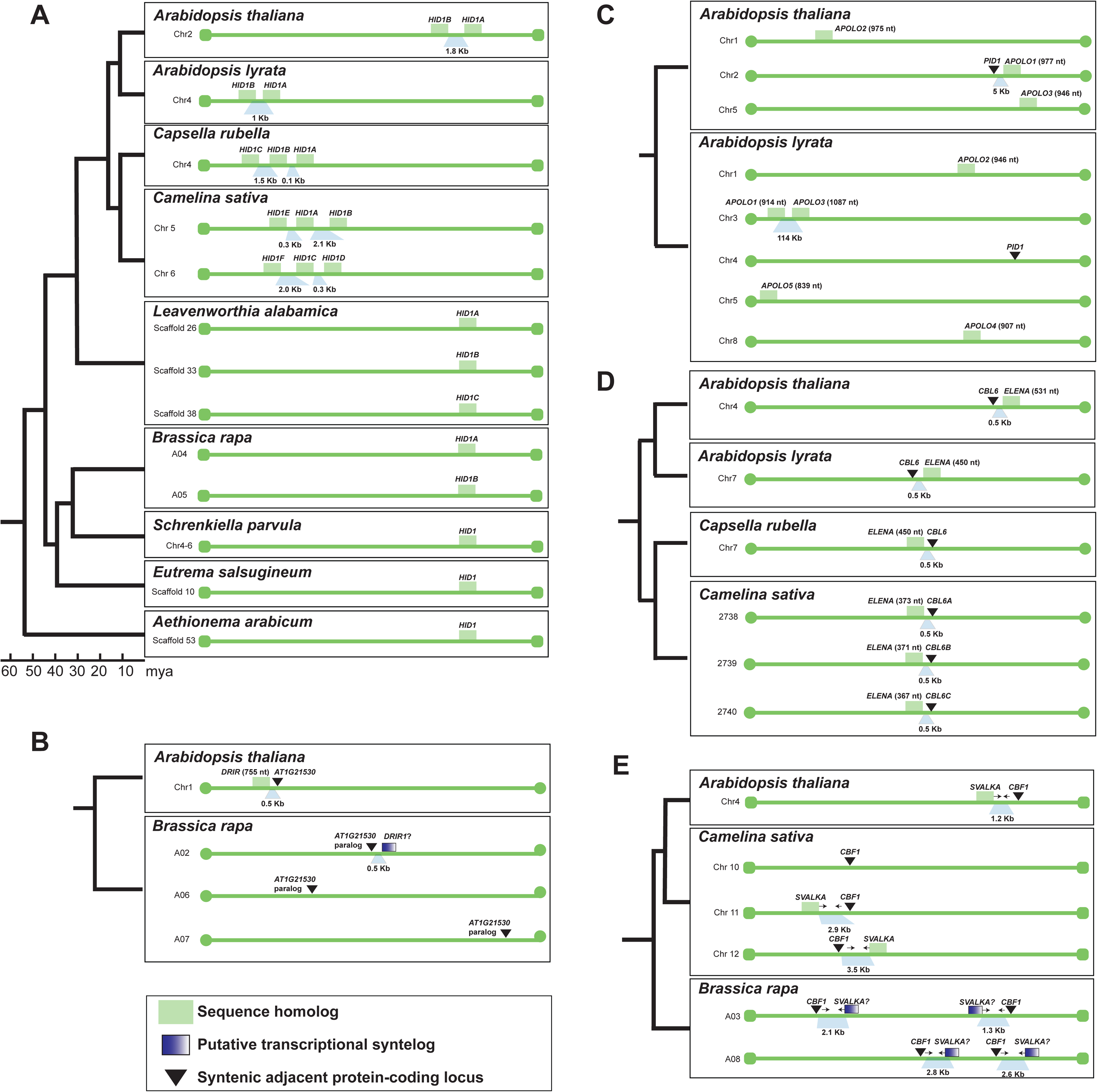
Evolution of functionally characterized lincRNAs. **A**) Schematic representing conservation of the HID1 locus across representative Brassicaceae. In Arabidopsis HID1A represents the published HID1 locus. All other green boxes represent loci inferred based on sequence homology and synteny. **B**) Conservation of DRIR1. Although no sequence homologs were identified for DRIR1, a putative transcriptional syntelog (Blue box) was identified in Brassica at a syntenic locus. **C**) Conservation of APOLO. Although APOLO sequence homologs (i.e., paralogs) were identified in Arabidopsis lyrata, none were adjacent to the PID1 ortholog, the protein-coding gene known to be regulated by APOLO in Arabidopsis thaliana. **D**) Conservation of ELENA. ELENA sequence homologs were identified in species as distantly related as Camelina sativa, where they were situated in syntenic positions adjacent to CBL6 orthologs. **E**) Conservation of SVALKA. Sequence homologs of SVALKA were identified in Camelina adjacent to CBF1. In Brassica, no sequence homologs were identified, but several putative transcriptional syntelogs were recovered adjacent to CBF1 orthologs.

**Supplemental Figure 8:**
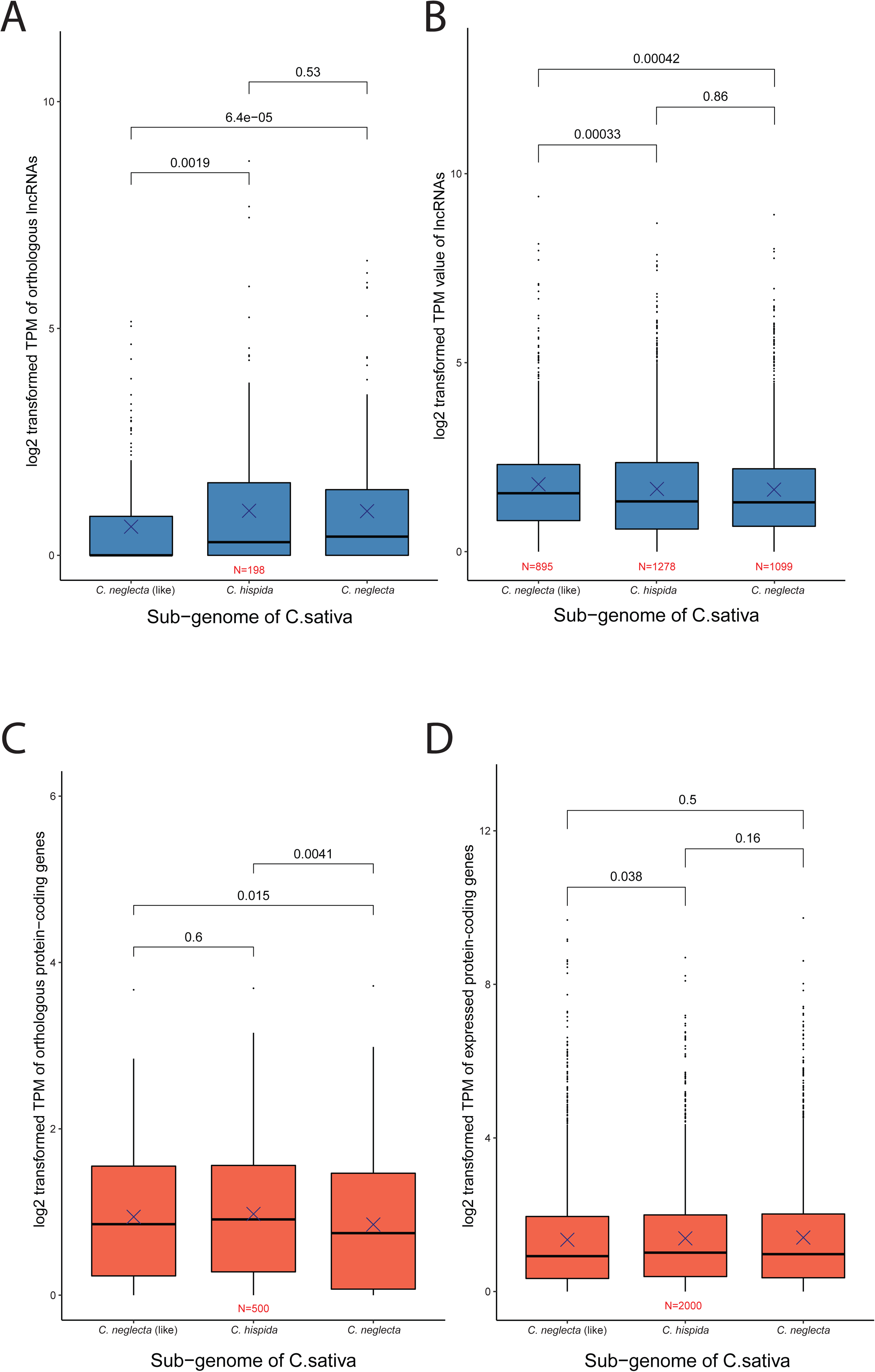
Subgenome expression dominance of *Camelina* lincRNAs. **A**) Expression values of lincRNAs found in all three subgenomes of *C. sativa.* **B**) Expression values of lincRNAs specific to one subgenome. **C**) Expression values of protein coding genes found in all three subgenomes of *C. sativa.* **D**) Expression values of protein coding genes specific to one subgenome. Numbers represent Student’s t-test *P* values between groups of expression values.

**Supplemental Figure 9:**
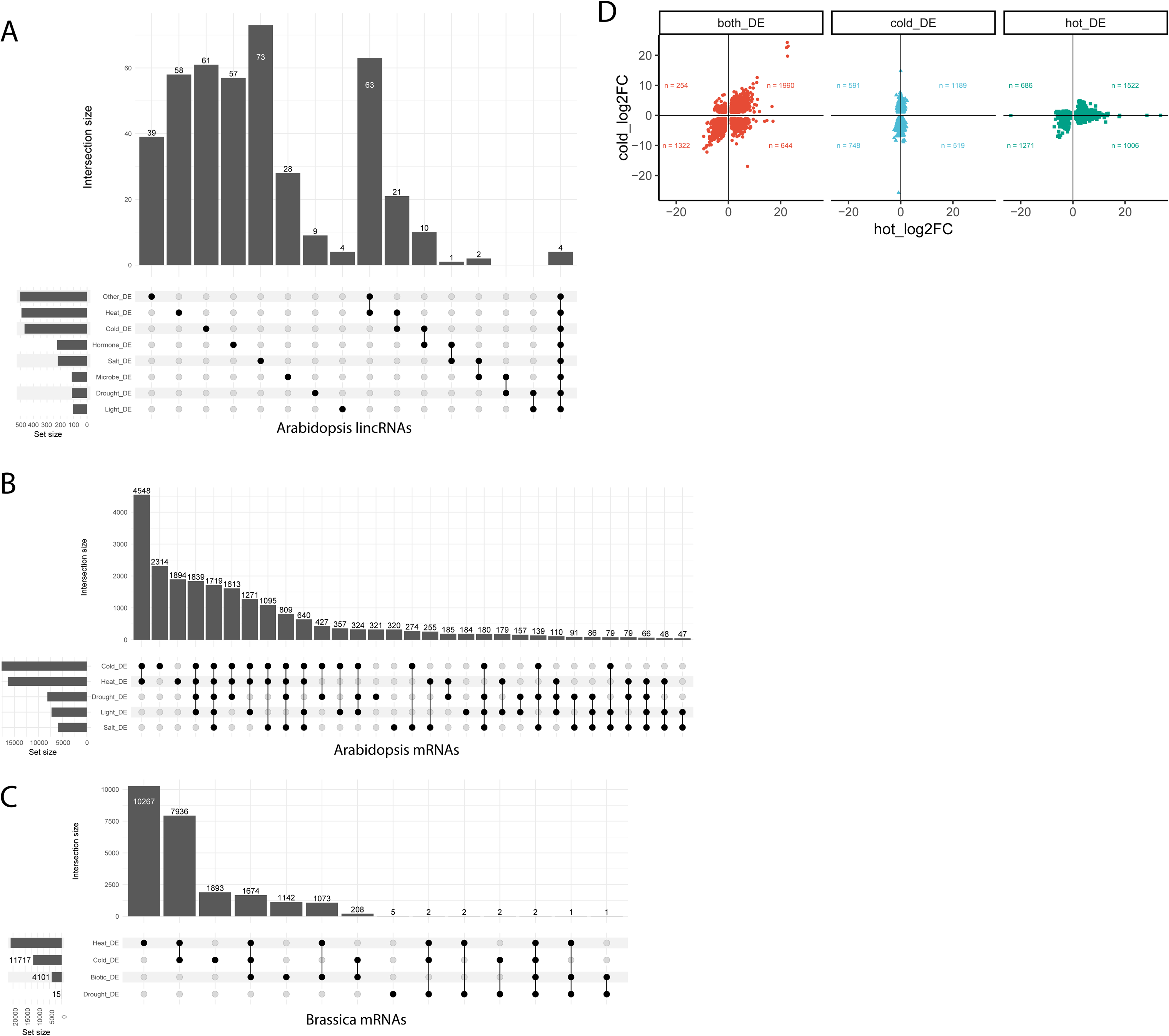
Differential expression during stress. **A**) Upset plot of Arabidopsis lincRNAs differentially expressed in a variety of broad stress categories (an expanded set of stresses compared to **Figure 6A**). **B**) Upset plot of Arabidopsis mRNAs found to be differentially expressed in various abiotic stresses. **C**) Upset plot of Brassica mRNAs found to be differentially expressed in various broad stress categories. **D**) Scatterplot comparing log2FC of Arabidopsis mRNAs in cold and heat stress when mRNAs are DE in both, or just a single stress. Note that a majority of differentially expressed mRNAs are up or down in both conditions.

**Supplemental Figure 10:**
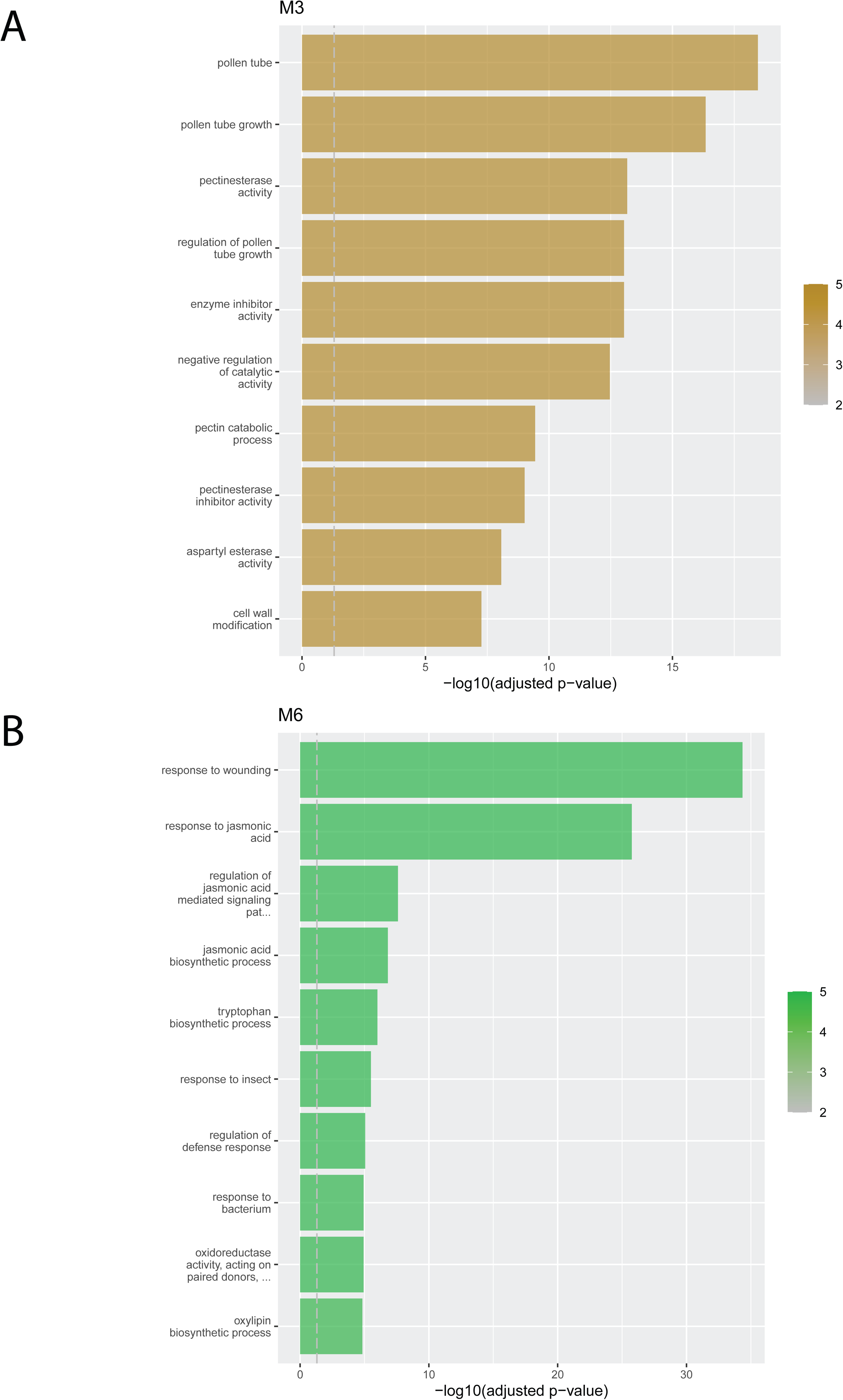
Enriched GO terms of mRNAs found in WGCNA modules from Figure 6E (**A**) and Figure 6F (**B**).

**Supplemental Figure 11:**
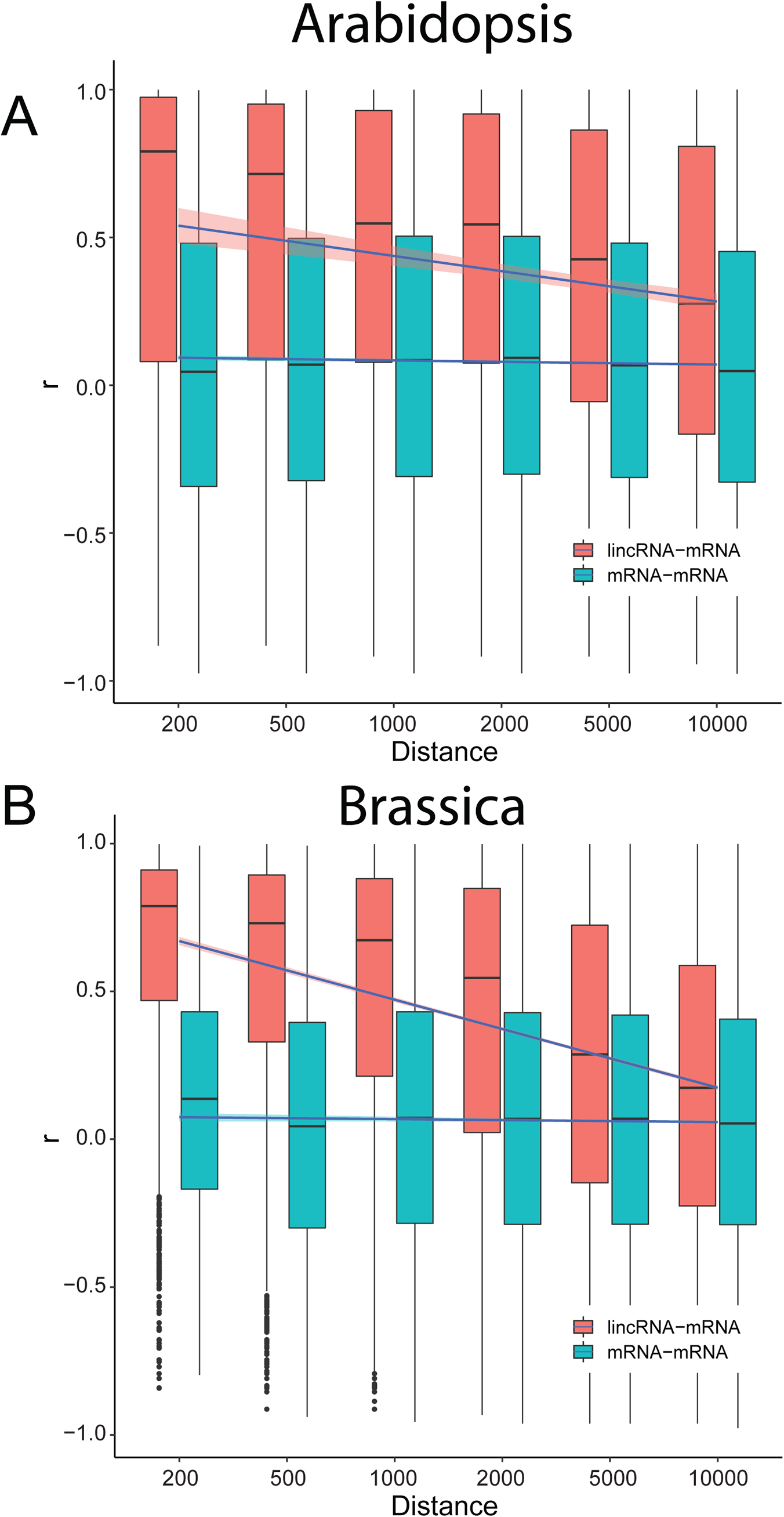
Gene expression correlation (Pearson) between lincRNA-mRNA and mRNA-mRNA pairs within defined distances in Arabidopsis and Brassica tissue atlases. **A**) Arabidopsis gene expression correlation of all expressed lincRNA/mRNAs with nearby expressed mRNAs within defined distances (x-axis). **B**) Brassica gene expression correlation of all expressed lincRNA/mRNAs with nearby expressed mRNAs within defined distances (x-axis). Note, all pairs within smaller distances are contained within larger distances.

**Supplemental Figure 12:**
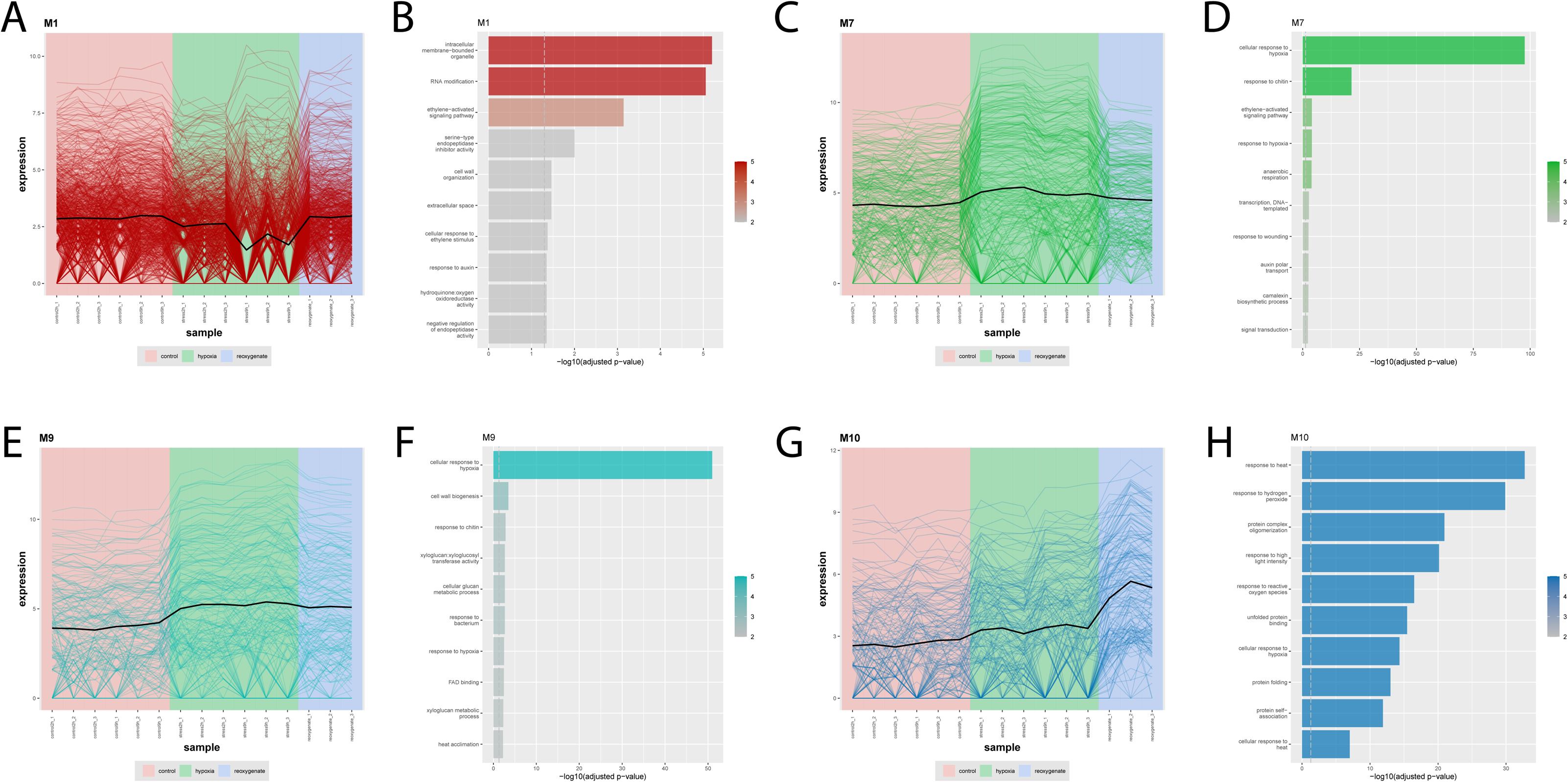
WGCNA modules of biological interest from the Lee and Serres hypoxia datasets containing lincRNAs. Expression profiles and enriched GO terms from module 1 (**A and B**), module 7 (**C and D**), module 9 (**E and F**), and module 10 (**G and H**).

**Supplemental File 1:** List of SRAs examined for all four species

**Supplemental File 2:** Functional annotations for each lincRNA from all four species

**Supplemental File 3:** Evolinc II results for all four species

**Supplemental File 4:** sORF and structural motif conservation and characteristics

**Supplemental File 5:** Predicted miRNA binding motifs for all four species

**Supplemental File 6:** SRAs, with associated metadata, used in targeted transcriptomic studies

**Supplemental File 7:** Araport11 lincRNAs that were removed from analysis

**Supplemental File 8:** Expression information for mRNAs and lincRNAs in paired analyses

## References

1. A Eccles, David. 2019. “Demultiplexing Nanopore Reads with LAST v6.” *Protocols.io*. ZappyLab, Inc. https://doi.org/10.17504/protocols.io.7vmhn46.

2. Amor, Besma Ben, Sonia Wirth, Francisco Merchan, Philippe Laporte, Yves d’Aubenton-Carafa, Judith Hirsch, Alexis Maizel, et al. 2009. “Novel Long Non-Protein Coding RNAs Involved in Arabidopsis Differentiation and Stress Responses.” Genome Research 19 (1): 57–69.

3. Ariel, Federico, Leandro Lucero, Aurelie Christ, Maria Florencia Mammarella, Teddy Jegu, Alaguraj Veluchamy, Kiruthiga Mariappan, et al. 2020. “R-Loop Mediated Trans Action of the APOLO Long Noncoding RNA.” Molecular Cell 77 (5): 1055–65.e4.

4. Bilichak, Andriy, Yaroslav Ilnytskyy, Rafal Wóycicki, Nina Kepeshchuk, Dawson Fogen, and Igor Kovalchuk. 2015. “The Elucidation of Stress Memory Inheritance in Brassica Rapa Plants.” Frontiers in Plant Science 6 (January): 5.

5. Bolstad, Ben. n.d. *preprocessCore*. Github. Accessed August 26, 2021. https://github.com/bmbolstad/preprocessCore.

6. Brandão, Marcelo M., Luiza L. Dantas, and Marcio C. Silva-Filho. 2009. “AtPIN: Arabidopsis Thaliana Protein Interaction Network.” BMC Bioinformatics 10 (December): 454.

7. Brown, C. J., B. D. Hendrich, J. L. Rupert, R. G. Lafrenière, Y. Xing, J. Lawrence, and H. F. Willard. 1992. “The Human XIST Gene: Analysis of a 17 Kb Inactive X-Specific RNA That Contains Conserved Repeats and Is Highly Localized within the Nucleus.” Cell 71 (3): 527–42.

8. Cabili, Moran N., Cole Trapnell, Loyal Goff, Magdalena Koziol, Barbara Tazon-Vega, Aviv Regev, and John L. Rinn. 2011. “Integrative Annotation of Human Large Intergenic Noncoding RNAs Reveals Global Properties and Specific Subclasses.” Genes & Development 25 (18): 1915–27.

9. Cantarel, Brandi L., Ian Korf, Sofia M. C. Robb, Genis Parra, Eric Ross, Barry Moore, Carson Holt, Alejandro Sánchez Alvarado, and Mark Yandell. 2008. “MAKER: An Easy-to-Use Annotation Pipeline Designed for Emerging Model Organism Genomes.” Genome Research 18 (1): 188–96.

10. Cheng, Shifeng, Erik van den Bergh, Peng Zeng, Xiao Zhong, Jiajia Xu, Xin Liu, Johannes Hofberger, et al. 2013. “The Tarenaya Hassleriana Genome Provides Insight into Reproductive Trait and Genome Evolution of Crucifers.” The Plant Cell 25 (8): 2813–30.

11. Chen, Ho-Ming, Yi-Hang Li, and Shu-Hsing Wu. 2007. “Bioinformatic Prediction and Experimental Validation of a microRNA-Directed Tandem Trans-Acting siRNA Cascade in Arabidopsis.” Proceedings of the National Academy of Sciences of the United States of America 104 (9): 3318–23.

12. Csorba, Tibor, Julia I. Questa, Qianwen Sun, and Caroline Dean. 2014. “Antisense COOLAIR Mediates the Coordinated Switching of Chromatin States at FLC during Vernalization.” Proceedings of the National Academy of Sciences of the United States of America 111 (45): 16160–65.

13. Dai, Xinbin, Zhaohong Zhuang, and Patrick Xuechun Zhao. 2018. “psRNATarget: A Plant Small RNA Target Analysis Server (2017 Release).” Nucleic Acids Research 46 (W1): W49–54.

14. De Smet, Riet, Keith L. Adams, Klaas Vandepoele, Marc C. E. Van Montagu, Steven Maere, and Yves Van de Peer. 2013. “Convergent Gene Loss Following Gene and Genome Duplications Creates Single-Copy Families in Flowering Plants.” Proceedings of the National Academy of Sciences of the United States of America 110 (8): 2898–2903.

15. Dew-Budd, Kelly, Julie Cheung, Kyle Palos, Evan S. Forsythe, and Mark A. Beilstein. 2020. “Evolutionary and Biochemical Analyses Reveal Conservation of the Brassicaceae Telomerase Ribonucleoprotein Complex.” PloS One 15 (4): e0222687.

16. Domon, Bruno, and Ruedi Aebersold. 2006. “Mass Spectrometry and Protein Analysis.” Science 312 (5771): 212–17.

17. Durinck, Steffen, Yves Moreau, Arek Kasprzyk, Sean Davis, Bart De Moor, Alvis Brazma, and Wolfgang Huber. 2005. “BioMart and Bioconductor: A Powerful Link between Biological Databases and Microarray Data Analysis.” Bioinformatics 21 (16): 3439–40.

18. ENCODE Project Consortium. 2012. “An Integrated Encyclopedia of DNA Elements in the Human Genome.” Nature 489 (7414): 57–74.

19. Fajkus, Petr, Vratislav Peška, Michal Závodník, Miloslava Fojtová, Jana Fulnečková, Šimon Dobias, Agata Kilar, et al. 2019. “Telomerase RNAs in Land Plants.” Nucleic Acids Research 47 (18): 9842–56.

20. Feng, J., W. D. Funk, S. S. Wang, S. L. Weinrich, A. A. Avilion, C. P. Chiu, R. R. Adams, E. Chang, R. C. Allsopp, and J. Yu. 1995. “The RNA Component of Human Telomerase.” Science 269 (5228): 1236–41.

21. Fernandez, Nicolas F., Gregory W. Gundersen, Adeeb Rahman, Mark L. Grimes, Klarisa Rikova, Peter Hornbeck, and Avi Ma’ayan. 2017. “Clustergrammer, a Web-Based Heatmap Visualization and Analysis Tool for High-Dimensional Biological Data.” Scientific Data 4 (October): 170151.

22. Foley, Shawn W., Sager J. Gosai, Dongxue Wang, Nur Selamoglu, Amelia C. Sollitti, Tino Köster, Alexander Steffen, et al. 2017. “A Global View of RNA-Protein Interactions Identifies Post-Transcriptional Regulators of Root Hair Cell Fate.” Developmental Cell 41 (2): 204–20.e5.

23. Gil, Noa, and Igor Ulitsky. 2020. “Regulation of Gene Expression by Cis-Acting Long Non-Coding RNAs.” Nature Reviews. Genetics 21 (2): 102–17.

24. Gosai, Sager J., Shawn W. Foley, Dongxue Wang, Ian M. Silverman, Nur Selamoglu, Andrew D. L. Nelson, Mark A. Beilstein, Fevzi Daldal, Roger B. Deal, and Brian D. Gregory. 2015. “Global Analysis of the RNA-Protein Interaction and RNA Secondary Structure Landscapes of the Arabidopsis Nucleus.” Molecular Cell 57 (2): 376–88.

25. Haug-Baltzell, Asher, Sean A. Stephens, Sean Davey, Carlos E. Scheidegger, and Eric Lyons. 2017. “SynMap2 and SynMap3D: Web-Based Whole-Genome Synteny Browsers.” Bioinformatics 33 (14): 2197–98.

26. Heinz, Sven, Christopher Benner, Nathanael Spann, Eric Bertolino, Yin C. Lin, Peter Laslo, Jason X. Cheng, Cornelis Murre, Harinder Singh, and Christopher K. Glass. 2010. “Simple Combinations of Lineage-Determining Transcription Factors Prime Cis-Regulatory Elements Required for Macrophage and B Cell Identities.” Molecular Cell 38 (4): 576–89.

27. Hong, Seong Hyeon, Jun Tae Kwon, Jihye Kim, Juri Jeong, Jaehwan Kim, Seonhee Lee, and Chunghee Cho. 2018. “Profiling of Testis-Specific Long Noncoding RNAs in Mice.” BMC Genomics 19 (1): 539.

28. Howe, Kevin L., Premanand Achuthan, James Allen, Jamie Allen, Jorge Alvarez-Jarreta, M. Ridwan Amode, Irina M. Armean, et al. 2021. “Ensembl 2021.” Nucleic Acids Research 49 (D1): D884–91.

29. Howell, Miya D., Noah Fahlgren, Elisabeth J. Chapman, Jason S. Cumbie, Christopher M. Sullivan, Scott A. Givan, Kristin D. Kasschau, and James C. Carrington. 2007. “Genome-Wide Analysis of the RNA-DEPENDENT RNA POLYMERASE6/DICER-LIKE4 Pathway in Arabidopsis Reveals Dependency on miRNA-and tasiRNA-Directed Targeting.” The Plant Cell 19 (3): 926–42.

30. Hsu, Polly Yingshan, Lorenzo Calviello, Hsin-Yen Larry Wu, Fay-Wei Li, Carl J. Rothfels, Uwe Ohler, and Philip N. Benfey. 2016. “Super-Resolution Ribosome Profiling Reveals Unannotated Translation Events in Arabidopsis.” Proceedings of the National Academy of Sciences. https://doi.org/10.1073/pnas.1614788113.

31. Ingolia, Nicholas T., Sina Ghaemmaghami, John R. S. Newman, and Jonathan S. Weissman. 2009. “Genome-Wide Analysis in Vivo of Translation with Nucleotide Resolution Using Ribosome Profiling.” Science 324 (5924): 218–23.

32. Ji, Zhe, Ruisheng Song, Aviv Regev, and Kevin Struhl. 2015. “Many lncRNAs, 5’UTRs, and Pseudogenes Are Translated and Some Are Likely to Express Functional Proteins.” eLife 4 (December): e08890.

33. Khyzha, Nadiya, Melvin Khor, Peter V. DiStefano, Liangxi Wang, Ljubica Matic, Ulf Hedin, Michael D. Wilson, Lars Maegdefessel, and Jason E. Fish. 2019. “Regulation of CCL2 Expression in Human Vascular Endothelial Cells by a Neighboring Divergently Transcribed Long Noncoding RNA.” Proceedings of the National Academy of Sciences of the United States of America 116 (33): 16410–19.

34. Kindgren, Peter, Ryan Ard, Maxim Ivanov, and Sebastian Marquardt. 2019. “Author Correction: Transcriptional Read-through of the Long Non-Coding RNA SVALKA Governs Plant Cold Acclimation.” Nature Communications 10 (1): 5141.

35. Klepikova, Anna V., Artem S. Kasianov, Evgeny S. Gerasimov, Maria D. Logacheva, and Aleksey A. Penin. 2016. “A High Resolution Map of the Arabidopsis Thaliana Developmental Transcriptome Based on RNA-Seq Profiling.” The Plant Journal: For Cell and Molecular Biology 88 (6): 1058–70.

36. Kopp, Florian, and Joshua T. Mendell. 2018. “Functional Classification and Experimental Dissection of Long Noncoding RNAs.” Cell 172 (3): 393–407.

37. Kovaka, Sam, Aleksey V. Zimin, Geo M. Pertea, Roham Razaghi, Steven L. Salzberg, and Mihaela Pertea. 2019. “Transcriptome Assembly from Long-Read RNA-Seq Alignments with StringTie2.” Genome Biology 20 (1): 278.

38. Krueger, Felix, and Simon R. Andrews. 2011. “Bismark: A Flexible Aligner and Methylation Caller for Bisulfite-Seq Applications.” Bioinformatics 27 (11): 1571–72.

39. Kuhn, Max, and Hadley Wickham. 2020. “Tidymodels: A Collection of Packages for Modeling and Machine Learning Using Tidyverse Principles.” Boston, MA, USA. [(accessed on 10 December 2020)].

40. Lamesch, Philippe, Tanya Z. Berardini, Donghui Li, David Swarbreck, Christopher Wilks, Rajkumar Sasidharan, Robert Muller, et al. 2012. “The Arabidopsis Information Resource (TAIR): Improved Gene Annotation and New Tools.” Nucleic Acids Research 40 (Database issue): D1202–10.

41. Lawrence, Michael, Wolfgang Huber, Hervé Pagès, Patrick Aboyoun, Marc Carlson, Robert Gentleman, Martin T. Morgan, and Vincent J. Carey. 2013. “Software for Computing and Annotating Genomic Ranges.” PLoS Computational Biology 9 (8): e1003118.

42. Li, Heng. 2013. “Aligning Sequence Reads, Clone Sequences and Assembly Contigs with BWA-MEM.” arXiv [q-bio.GN]. arXiv. http://arxiv.org/abs/1303.3997.

43. Li, Heng. 2018. “Minimap2: Pairwise Alignment for Nucleotide Sequences.” Bioinformatics 34 (18): 3094–3100.

44. Li, Lin, Steven R. Eichten, Rena Shimizu, Katherine Petsch, Cheng-Ting Yeh, Wei Wu, Antony M. Chettoor, et al. 2014. “Genome-Wide Discovery and Characterization of Maize Long Non-Coding RNAs.” Genome Biology 15 (2): R40.

45. Liu, Jun, Choonkyun Jung, Jun Xu, Huan Wang, Shulin Deng, Lucia Bernad, Catalina Arenas-Huertero, and Nam-Hai Chua. 2012. “Genome-Wide Analysis Uncovers Regulation of Long Intergenic Noncoding RNAs in Arabidopsis.” The Plant Cell 24 (11): 4333–45.

46. Lorenzi, Lucia, Hua-Sheng Chiu, Francisco Avila Cobos, Stephen Gross, Pieter-Jan Volders, Robrecht Cannoodt, Justine Nuytens, et al. 2021. “The RNA Atlas Expands the Catalog of Human Non-Coding RNAs.” Nature Biotechnology, June. https://doi.org/10.1038/s41587-021-00936-1.

47. Love, Michael I., Wolfgang Huber, and Simon Anders. 2014. “Moderated Estimation of Fold Change and Dispersion for RNA-Seq Data with DESeq2.” Genome Biology 15 (12): 550.

48. Merchant, Nirav, Eric Lyons, Stephen Goff, Matthew Vaughn, Doreen Ware, David Micklos, and Parker Antin. 2016. “The iPlant Collaborative: Cyberinfrastructure for Enabling Data to Discovery for the Life Sciences.” PLoS Biology 14 (1): e1002342.

49. Moghe, Gaurav D., Melissa D. Lehti-Shiu, Alex E. Seddon, Shan Yin, Yani Chen, Piyada Juntawong, Federica Brandizzi, Julia Bailey-Serres, and Shin-Han Shiu. 2013. “Characteristics and Significance of Intergenic Polyadenylated RNA Transcription in Arabidopsis.” Plant Physiology 161 (1): 210–24.

50. Nelson, Andrew D. L., Upendra K. Devisetty, Kyle Palos, Asher K. Haug-Baltzell, Eric Lyons, and Mark A. Beilstein. 2017. “Evolinc: A Tool for the Identification and Evolutionary Comparison of Long Intergenic Non-Coding RNAs.” Frontiers in Genetics 8 (May): 52.

51. Neph, Shane, M. Scott Kuehn, Alex P. Reynolds, Eric Haugen, Robert E. Thurman, Audra K. Johnson, Eric Rynes, et al. 2012. “BEDOPS: High-Performance Genomic Feature Operations.” Bioinformatics 28 (14): 1919–20.

52. Patro, Rob, Geet Duggal, Michael I. Love, Rafael A. Irizarry, and Carl Kingsford. 2017. “Salmon Provides Fast and Bias-Aware Quantification of Transcript Expression.” Nature Methods 14 (4): 417–19.

53. Peri, Sateesh, Sarah Roberts, Isabella R. Kreko, Lauren B. McHan, Alexandra Naron, Archana Ram, Rebecca L. Murphy, et al. 2019. “Read Mapping and Transcript Assembly: A Scalable and High-Throughput Workflow for the Processing and Analysis of Ribonucleic Acid Sequencing Data.” Frontiers in Genetics 10: 1361.

54. Provart, Nicholas, and Tong Zhu. 2003. “A Browser-Based Functional Classification SuperViewer for Arabidopsis Genomics.” Curr Comput Mol Biol 2003: 271–72.

55. Qin, Tao, Huayan Zhao, Peng Cui, Nour Albesher, and Liming Xiong. 2017. “A Nucleus-Localized Long Non-Coding RNA Enhances Drought and Salt Stress Tolerance.” Plant Physiology 175 (3): 1321–36.

56. Qi, Xin, Shaojun Xie, Yuwei Liu, Fei Yi, and Jingjuan Yu. 2013. “Genome-Wide Annotation of Genes and Noncoding RNAs of Foxtail Millet in Response to Simulated Drought Stress by Deep Sequencing.” Plant Molecular Biology 83 (4-5): 459–73.

57. Quinlan, Aaron R., and Ira M. Hall. 2010. “BEDTools: A Flexible Suite of Utilities for Comparing Genomic Features.” Bioinformatics 26 (6): 841–42.

58. Ramírez, Fidel, Friederike Dündar, Sarah Diehl, Björn A. Grüning, and Thomas Manke. 2014. “deepTools: A Flexible Platform for Exploring Deep-Sequencing Data.” Nucleic Acids Research 42 (Web Server issue): W187–91.

59. Russo, Pedro S. T., Gustavo R. Ferreira, Lucas E. Cardozo, Matheus C. Bürger, Raul Arias-Carrasco, Sandra R. Maruyama, Thiago D. C. Hirata, et al. 2018. “CEMiTool: A Bioconductor Package for Performing Comprehensive Modular Co-Expression Analyses.” BMC Bioinformatics 19 (1): 56.

60. Seki, Masahide, Eri Katsumata, Ayako Suzuki, Sarun Sereewattanawoot, Yoshitaka Sakamoto, Junko Mizushima-Sugano, Sumio Sugano, et al. 2019. “Evaluation and Application of RNA-Seq by MinION.” DNA Research: An International Journal for Rapid Publication of Reports on Genes and Genomes 26 (1): 55–65.

61. Seo, Jun Sung, Hai-Xi Sun, Bong Soo Park, Chung-Hao Huang, Shyi-Dong Yeh, Choonkyun Jung, and Nam-Hai Chua. 2017. “ELF18-INDUCED LONG-NONCODING RNA Associates with Mediator to Enhance Expression of Innate Immune Response Genes in Arabidopsis.” The Plant Cell 29 (5): 1024–38.

62. Shuai, Peng, Dan Liang, Sha Tang, Zhoujia Zhang, Chu-Yu Ye, Yanyan Su, Xinli Xia, and Weilun Yin. 2014. “Genome-Wide Identification and Functional Prediction of Novel and Drought-Responsive lincRNAs in Populus Trichocarpa.” Journal of Experimental Botany 65 (17): 4975–83.

63. Soneson, Charlotte, Michael I. Love, and Mark D. Robinson. 2015. “Differential Analyses for RNA-Seq: Transcript-Level Estimates Improve Gene-Level Inferences.” F1000Research 4 (December): 1521.

64. Song, Jiarui, Dhenugen Logeswaran, Claudia Castillo-González, Yang Li, Sreyashree Bose, Behailu Birhanu Aklilu, Zeyang Ma, Alexander Polkhovskiy, Julian J-L Chen, and Dorothy E. Shippen. 2019. “The Conserved Structure of Plant Telomerase RNA Provides the Missing Link for an Evolutionary Pathway from Ciliates to Humans.” Proceedings of the National Academy of Sciences of the United States of America 116 (49): 24542–50.

65. Sun, Yutong, and Li Ma. 2019. “New Insights into Long Non-Coding RNA MALAT1 in Cancer and Metastasis.” Cancers 11 (2). https://doi.org/10.3390/cancers11020216.

66. Team, R. Core, and Others. 2013. “R: A Language and Environment for Statistical Computing.” http://r.meteo.uni.wroc.pl/web/packages/dplR/vignettes/intro-dplR.pdf.

67. Tian, Weidong, Lan V. Zhang, Murat Taşan, Francis D. Gibbons, Oliver D. King, Julie Park, Zeba Wunderlich, J. Michael Cherry, and Frederick P. Roth. 2008. “Combining Guilt-by-Association and Guilt-by-Profiling to Predict Saccharomyces Cerevisiae Gene Function.” Genome Biology 9 Suppl 1 (June): S7.

68. Tong, Chaobo, Xiaowu Wang, Jingyin Yu, Jian Wu, Wanshun Li, Junyan Huang, Caihua Dong, Wei Hua, and Shengyi Liu. 2013. “Comprehensive Analysis of RNA-Seq Data Reveals the Complexity of the Transcriptome in Brassica Rapa.” BMC Genomics 14 (October): 689.

69. Wang, Jia, Li Song, Xue Gong, Jinfan Xu, and Minhui Li. 2020. “Functions of Jasmonic Acid in Plant Regulation and Response to Abiotic Stress.” International Journal of Molecular Sciences 21 (4). https://doi.org/10.3390/ijms21041446.

70. Wang, Yuqiu, Xiuduo Fan, Fang Lin, Guangming He, William Terzaghi, Danmeng Zhu, and Xing Wang Deng. 2014. “Arabidopsis Noncoding RNA Mediates Control of Photomorphogenesis by Red Light.” Proceedings of the National Academy of Sciences of the United States of America 111 (28): 10359–64.

71. West, Jason A., Christopher P. Davis, Hongjae Sunwoo, Matthew D. Simon, Ruslan I. Sadreyev, Peggy I. Wang, Michael Y. Tolstorukov, and Robert E. Kingston. 2014. “The Long Noncoding RNAs NEAT1 and MALAT1 Bind Active Chromatin Sites.” Molecular Cell 55 (5): 791–802.

72. Wu, Hsin-Yen Larry, and Polly Yingshan Hsu. 2021. “Actively Translated uORFs Reduce Translation and mRNA Stability Independent of NMD in Arabidopsis.” bioRxiv. https://doi.org/10.1101/2021.09.16.460672.

73. Wu, Hsin-Yen Larry, Gaoyuan Song, Justin W. Walley, and Polly Yingshan Hsu. 2019. “The Tomato Translational Landscape Revealed by Transcriptome Assembly and Ribosome Profiling.” Plant Physiology 181 (1): 367–80.

74. Yanai, Itai, Hila Benjamin, Michael Shmoish, Vered Chalifa-Caspi, Maxim Shklar, Ron Ophir, Arren Bar-Even, et al. 2005. “Genome-Wide Midrange Transcription Profiles Reveal Expression Level Relationships in Human Tissue Specification.” Bioinformatics 21 (5): 650–59.

75. Yang, Ruolin, David E. Jarvis, Hao Chen, Mark A. Beilstein, Jane Grimwood, Jerry Jenkins, Shengqiang Shu, et al. 2013. “The Reference Genome of the Halophytic Plant Eutrema Salsugineum.” Frontiers in Plant Science 4 (March): 46.

76. Zhang, Xiaona, Yanchun Zhou, Shaoying Chen, Wei Li, Weibing Chen, and Wei Gu. 2019. “LncRNA MACC1-AS1 Sponges Multiple miRNAs and RNA-Binding Protein PTBP1.” Oncogenesis 8 (12): 73.

77. Zhao, Xinyue, Jingrui Li, Bi Lian, Hanqing Gu, Yan Li, and Yijun Qi. 2018. “Global Identification of Arabidopsis lncRNAs Reveals the Regulation of MAF4 by a Natural Antisense RNA.” Nature Communications 9 (1): 5056.

